# Combining LIANA and Tensor-cell2cell to decipher cell-cell communication across multiple samples

**DOI:** 10.1101/2023.04.28.538731

**Authors:** Hratch Baghdassarian, Daniel Dimitrov, Erick Armingol, Julio Saez-Rodriguez, Nathan E. Lewis

## Abstract

In recent years, data-driven inference of cell-cell communication has helped reveal coordinated biological processes across cell types. While multiple cell-cell communication tools exist, results are specific to the tool of choice, due to the diverse assumptions made across computational frameworks. Moreover, tools are often limited to analyzing single samples or to performing pairwise comparisons. As experimental design complexity and sample numbers continue to increase in single-cell datasets, so does the need for generalizable methods to decipher cell-cell communication in such scenarios. Here, we integrate two tools, LIANA and Tensor-cell2cell, which combined can deploy multiple existing methods and resources, to enable the robust and flexible identification of cell-cell communication programs across multiple samples. In this protocol, we show how the integration of our tools facilitates the choice of method to infer cell-cell communication and subsequently perform an unsupervised deconvolution to obtain and summarize biological insights. We explain how to perform the analysis step-by-step in both Python and R, and we provide online tutorials with detailed instructions available at https://ccc-protocols.readthedocs.io/. This protocol typically takes ∼1.5h to complete from installation to downstream visualizations on a GPU-enabled computer, for a dataset of ∼63k cells, 10 cell types, and 12 samples.

## Introduction

Cell-cell communication (CCC) coordinates higher-order biological functions in multicellular organisms^1, 2^, dictating phenotypes in response to different contexts such as disease state, spatial location, and organismal life stage. In recent years, many tools have been developed to leverage single-cell and spatial transcriptomics data to understand CCC events driving various biological processes^2^. While each computational strategy contributes unique and valuable developments, many are tool-specific and challenging to integrate due to a plethora of different inference methods and resources housing prior knowledge^2, 3^. Moreover, most tools do not account for the relationships of coordinated CCC events (CCC programs) across different contexts^4^, either disregarding context altogether by analyzing samples individually or being limited to pairwise comparisons. Thus, as the ability to generate large single-cell and spatial transcriptomics datasets and the interest in studying CCC programs continues to increase^5–7^, the need to robustly decipher CCC is becoming essential.

### Development of the protocol

We combine two independent yet highly complementary tools that leverage existing methods to enable robust and hypothesis-free analysis of context-driven cell-cell communication programs (**Fig.1**). LIANA^3^ is a computational framework that implements multiple available ligand-receptor resources (i.e., database of ligand-receptor interactions) and methods to analyze CCC. In particular, the user can employ LIANA to select any method and resource of choice or combine multiple approaches simultaneously to obtain consensus predictions. Tensor-cell2cell^8^ is a dimensionality reduction approach devised to uncover context-driven CCC programs across multiple samples simultaneously. Specifically, Tensor-cell2cell uses CCC scores inferred by any method and arranges the data into a 4D tensor to capture the coordinated relationship between ligand-receptor interactions, communicating cell type pairs, and samples. Together, LIANA and Tensor-cell2cell unify existing approaches to enable researchers to easily use their preferred CCC resource and method and subsequently analyze any number of samples into biologically-relevant CCC insights without the additional complications of installing separate tools or reconciling discrepancies between them.

**Figure 1.**
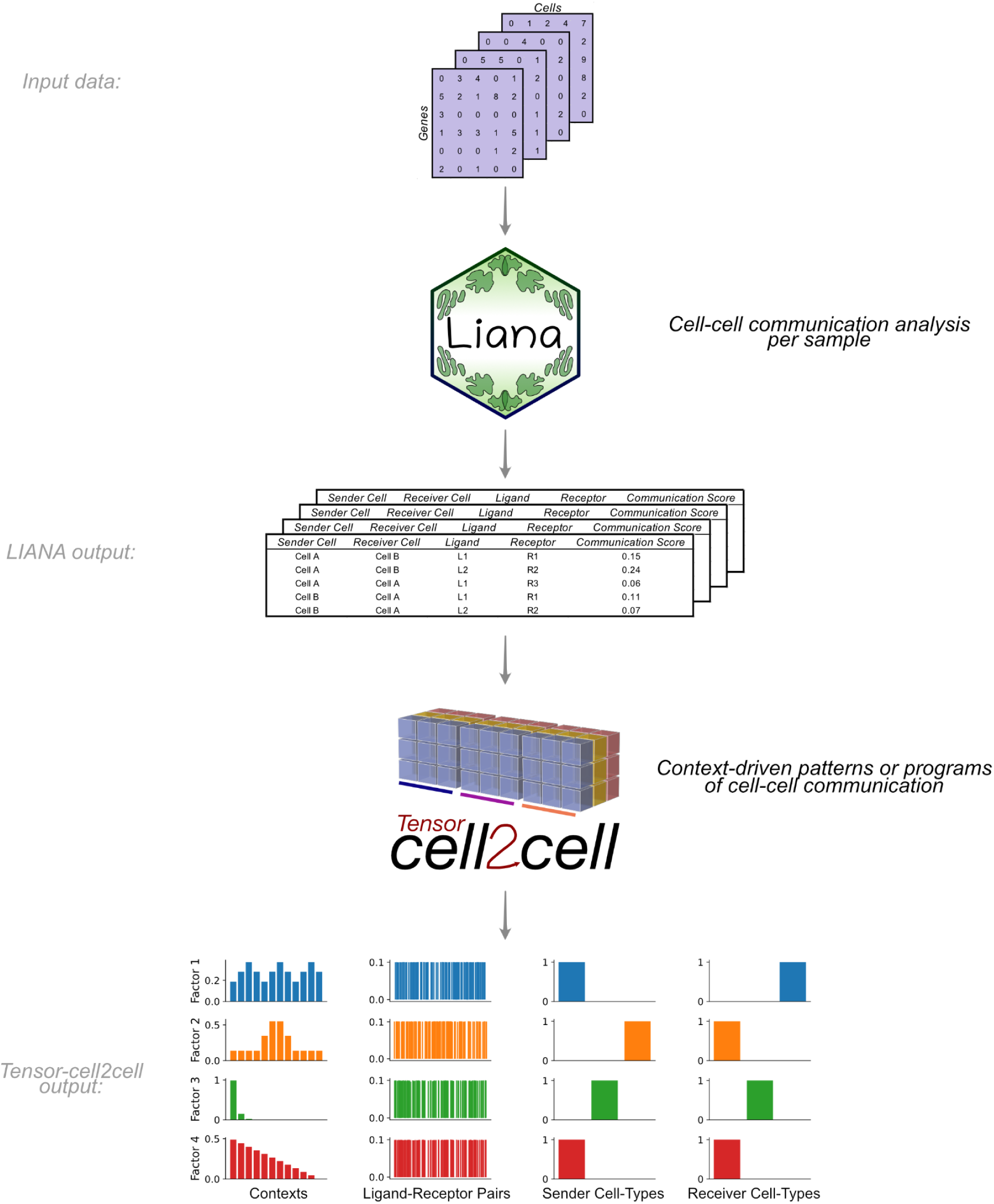
Integration of LIANA and Tensor-cell2cell to identify context-driven programs of cell-cell communication. LIANA and Tensor-cell2cell can be used together to infer the molecular basis of cell-cell interactions by running analysis across multiple samples, conditions or contexts. Given a method, resource, and expression data, LIANA outputs CCC scores for all interactions in a sample. We adapted both tools to be highly compatible with each other, so LIANA outputs can be directly passed to Tensor-cell2cell to detect the programs from the scores computed with LIANA. Tensor-cell2cell uses the communication scores generated for multiple samples to identify context-driven CCC programs.

For this protocol, we adapted LIANA and Tensor-cell2cell to enable their smooth integration. Thus, our protocol demonstrates the concerted use of both tools, describes the insights they provide, and facilitates the interpretation of their outputs. We base this protocol on recent best practices for single-cell transcriptomics and CCC inference^9^. We begin by processing the key inputs of our tools. Then, we guide the selection of methods and prior-knowledge resources to score intercellular communication, using LIANA’s consensus method and resource to infer the potential CCC events for each sample. We use Tensor-cell2cell to summarize the intercellular communication events across samples, and we describe key technical considerations to enable consistent decomposition results. Finally, we guide the interpretation of the decomposition results, and show multiple downstream analyses and visualizations to facilitate interpretation of the context-dependent CCC programs. For example, we illustrate how biologically-relevant results can be obtained by coupling the outputs with pathway-enrichment analyses. We also provide quickstart and in-depth online tutorials with detailed descriptions of all steps described in this protocol and their crucial parameters. All these materials are available in both Python and R at https://ccc-protocols.readthedocs.io/. Collectively, these materials provide a comprehensive and flexible playbook to investigate cell-cell communication from single-cell transcriptomics.

### Applications of the protocol

LIANA and Tensor-cell2cell have been used for diverse purposes. LIANA was initially used to compare and evaluate different ligand-receptor methods in diverse biological contexts. Tensor-cell2cell was originally applied to link CCC programs with different severities of COVID-19 and Autism Spectrum Disorder (ASD)^8^. Briefly, LIANA evaluated different methods and showed that they have limited agreement in terms of communication mechanisms^3, 8^, while Tensor-cell2cell revealed distinct CCC program dysregulations associated with severe COVID-19 specifically rather than moderate cases, as well as combinations of programs distinguishing ASD from neurotypical condition. Notably, LIANA provides a consensus resource and can aggregate multiple methods into consensus communication scores. Additionally, there is a natural complementarity between the two tools, as Tensor-cell2cell can use input scores from any CCC method (**Fig.1**) and generates consistent decomposition results across methods. Thus, our tools are highly generalizable and applicable to the analysis of any single-cell transcriptomics datasets. For example, LIANA has been used for the analysis of myocardial infarction^10^ and TGFβ signaling in breast cancer^11^, among others. Our tools are also applicable to other data modalities containing potentially interacting cell populations. Specifically, one can adapt LIANA or use existing spatial tools^12^ and combine their outputs with Tensor-cell2cell to generate spatially-informed CCC insights across contexts. Similarly, one can also obtain metabolite-mediated intercellular interactions^13, 14^, and decompose those into patterns across contexts with Tensor-cell2cell^15^. One can also apply Tensor-cell2cell to extract CCC programs occurring at specific tissues^16^ or at a whole-body organism level^16, 17^. In this protocol, we focus on how one can leverage the different CCC methods and resources, generalized by LIANA, to infer context-dependent CCC programs with Tensor-cell2cell from single-cell transcriptomics data.

### Comparison with other methods

A plethora of ligand-receptor methods have emerged, most of which were published with their own resources^1, 3, 8^. Many of these provide distinct scoring functions to prioritize interactions, yet studies have reported low agreement between their predictions^3, 18, 19^. Due to the lack of a gold standard, the benchmark of these methods remains limited^2, 3^ and it is challenging to choose the method that works best. To this end, in addition to providing multiple individual methods via LIANA, we also enable their consensus, which we use in this protocol, under the assumption that the wisdom of the crowd is less biased than any individual method^20^.

While many methods exist to infer ligand-receptor interactions from a single sample, fewer approaches were designed to compare CCC interactions across conditions. These include CrossTalkeR^21^, which utilizes network topological measures to compare communication patterns, CellPhoneDB^22^, which accepts user-provided lists of differentially-expressed genes to return relevant ligand-receptor interactions, and scDiffCom^23^, which uses a combined permutation approach across both cell types and conditions. Still, the aforementioned approaches are limited to pairwise comparisons. To our knowledge the only approach other than Tensor-cell2cell that can handle more than two conditions is CellChat^24^; however it is still based on pairwise comparisons, subsequently applying a manifold learning to summarize pathway-focused similarities of contexts. A key advantage of Tensor-cell2cell is that it considers all samples simultaneously while preserving the relationships between ligand-receptor interactions and communicating cell-type pairs. Thus, Tensor-cell2cell preserves higher-order CCC relationships and translates those into mechanistic CCC programs of potentially interacting ligands, receptors and communicating cell types.

### Limitations

Although our tools provide robust and flexible solutions to infer CCC patterns across contexts, they inherit the limitations associated with inferring communication events from transcriptomics data. These include the assumption that gene co-expression is indicative of active signaling events, which are largely mediated by proteins and their interactions, while also disregarding any biological processes, such as protein translation, post-translational modifications, secretion, diffusion, and trigger of intracellular events that precede and follow the interaction itself^2, 3^. Moreover, the aggregation of single cells into cell groups is essential when inferring potential CCC events, which could occlude some signals in heterogeneous tissues^2^, thereby biasing the insights that can be obtained. Finally, since the input of Tensor-cell2cell is a 4D-tensor, it requires that all elements are measured across all features and samples. Consequently, one should consider how to handle missing values caused by samples that do not present the same cell types and/or expressed genes when constructing this tensor. Deciding whether those reflect biologically-meaningful zeroes or a technical artifact may lead to variations in the resulting CCC patterns. We provide an discussion of the related parameter choices that may help users decide how to handle this challenge.

### Expertise needed to implement the protocol

Our protocol requires basic understanding of Python or R and single-cell data analysis. Yet, some of the detailed tutorials also touch on considerations that would be of interest to computational biologist power users.

## Materials

### Equipment

#### Hardware

This protocol was run on a computer with the following specifications:

● CPU: AMD Ryzen Threadripper 3960x (24 cores)
● Memory: 128GB DDR4
● GPU: NVIDIA RTX A6000 48GB

However, the minimal requirements for running this protocol are:

● CPU: 64-bit Intel or AMD processor (4 cores)
● Memory: 16GB DDR3
● GPU: NVIDIA GTX 1050 Ti (Optional)
● Storage: At least 10GB available

#### Software

#### Equipment setup

To facilitate the setup of a virtual environment containing all required packages with their corresponding versions, we provide an executable ‘setup_env.sh‘ script together with instructions on a Github repository we prepared for this protocol: https://github.com/saezlab/ccc_protocols/tree/main/env_setup

## Procedure

**Δ CRITICAL** In this section we introduce our protocol (**Fig.2**) using Python. The same protocol is implemented in R and is available online at https://ccc-protocols.readthedocs.io/en/latest/notebooks/ccc_R/QuickStart.html.

1. **Installation and Environment Setup**

**Figure 2.**
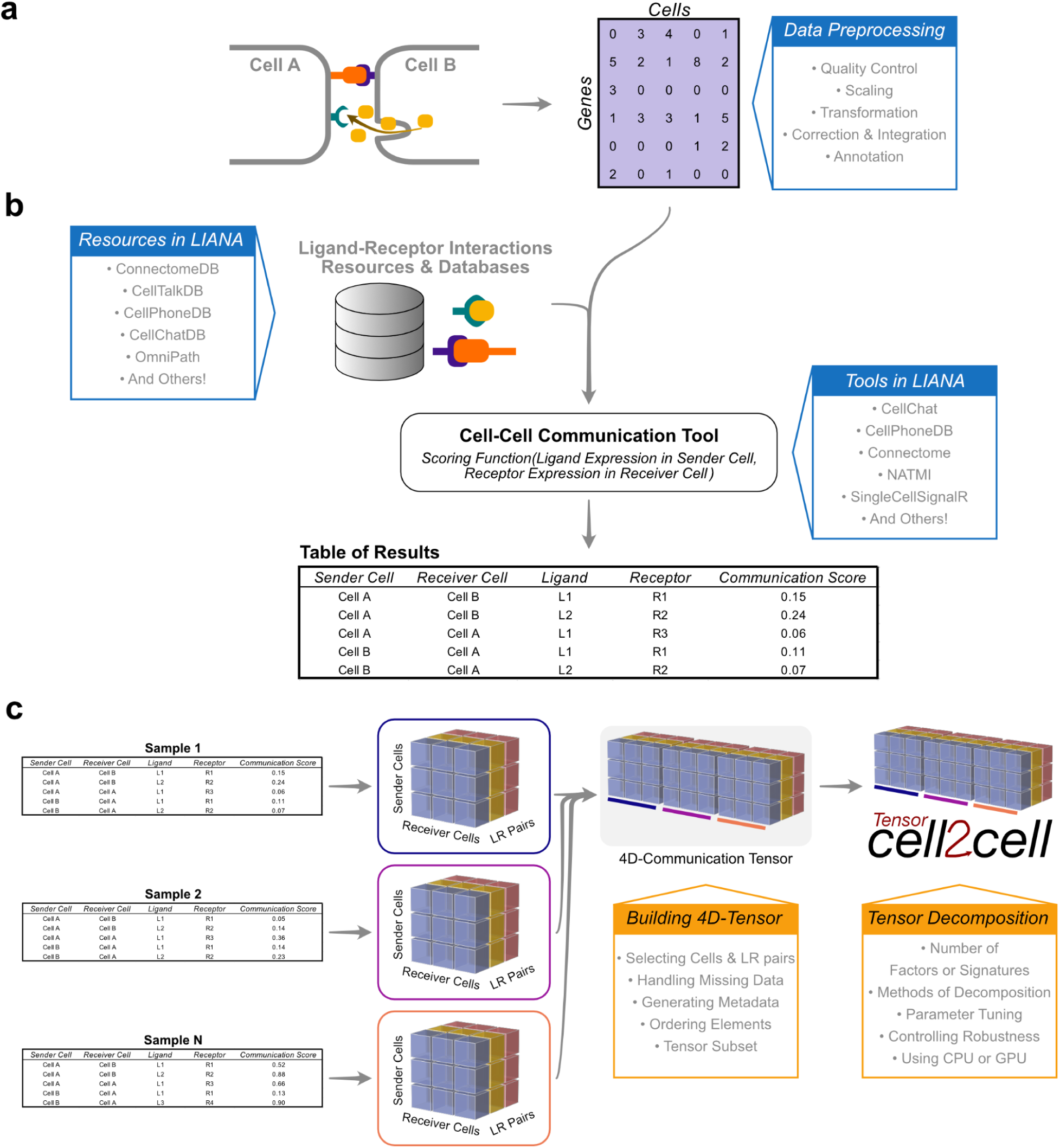
Overview of the protocol for inferring cell-cell communication through LIANA and Tensor-cell2cell. Main inputs, steps, resources and options are summarized for the distinct steps of this protocol: (**a**) A preprocessed gene expression matrix according to the best practices of single-cell analysis is expected as input (step 3 in the Procedure section). (**b**) This input data is integrated with the ligand-receptor resources available in LIANA to infer cell-cell communication using any of the methods implemented in LIANA (step 4 in the Procedure section). An output containing the cell-cell communication scores across all interactions per sample is generated. (**c**) The LIANA output is then directly passed to Tensor-cell2cell to build the respective communication tensor used by the tensor component analysis (steps 5.1-5.2 in the Procedure section). The output generated by Tensor-cell2cell can be later employed for other downstream analyses (steps 5.3 and 6 in the Procedure section).

Install Anaconda or Miniconda through the official instructions at: https://docs.anaconda.com/anaconda/install/index.html

Then, open a terminal to create and activate a conda environment:

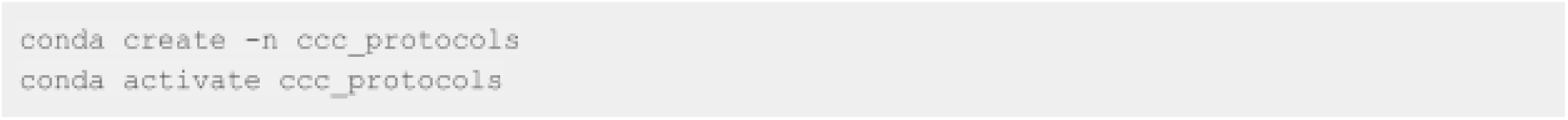

If you will be using a GPU, install PyTorch using conda:

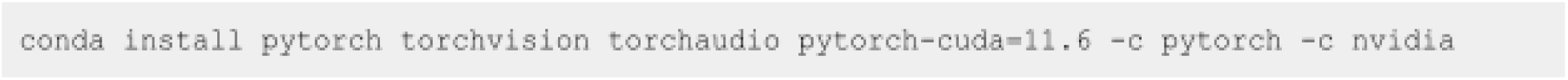

Install Tensor-cell2cell, LIANA, and decoupler using PyPI:

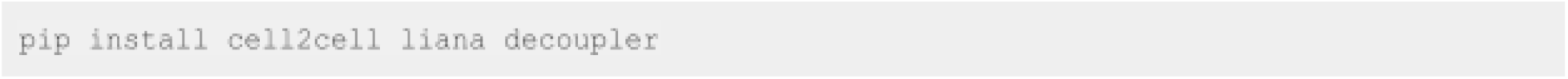

For fully reproducible runs of our Tutorials in both Python and R, we have specified the required packages and their versions in **Table 1**. You can also follow instructions in the *Environment setup* section to install a clean virtual environment with all package requirements.

**Table 1.** Required packages for the computational environment.

| Package Name | Package Version | Language | Install With |
| --- | --- | --- | --- |
| jupyter |  |  | conda |
| ipywidgets |  |  | conda |
| pip | >=22 | Python | conda |
| scanpy | >=1.9 | Python | conda |
| *cuda-toolkit |  |  | conda |
| *pytorch-cuda | 11.6 |  | conda |
| *torchvision |  |  | conda |
| *torchaudio |  |  | conda |
| pytorch, *cuda enabled |  |  | conda |
| scvi-tools | >=0.18 | Python | conda |
| scikit-misc | 0.1.4 | Python | conda |
| cell2cell | 0.6.7 | Python | pip |
| liana | 0.1.7 | Python | pip |
| decoupler | 1.3.3 | Python | pip |
| omnipath | 1.0.6 | Python | pip |
| singlecellexperiment |  | R | conda |
| remotes | >=2 | R | conda |
| devtools | >=2 | R | conda |
| seuratobject |  | R | conda |
| biocmanager | >=1.30 | R | conda |
| seurat | >=4 | R | conda |
| hd5r |  | R | conda |
| furrr |  | R | conda |
| textshape |  | R | conda |
| forcats |  | R | conda |
| rstatix |  | R | conda |
| ggpubr |  | R | conda |
| scater |  | R | conda |
| zellkonverter |  | R | conda |
| liana | Commit ID:<br>ab70b34066f68df60e9ed0d<br>0ce72b0d00f871b7e | R | remotes |
| seurat-disk | Commit ID:<br>9b89970eac2a3bd770e744f<br>63c7763419486b14c | R | remotes |
| decoupleR | Commit ID:<br>c17d635e0720c86f2386c39<br>ad7dea8614df393f1 | R | biocmanager |

\*: For GPU enabled use only
Python packages should always be installed. R language packages only need to be installed if planning to run the notebooks in R.

Notebooks to run this tutorial can be created by starting jupyter notebook:

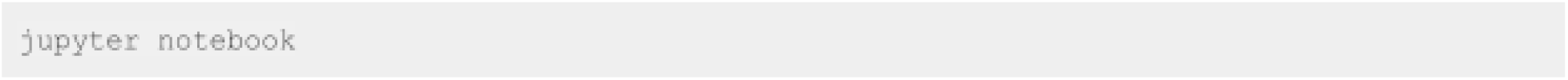

**2. Initial Setups**

First, if you are using a NVIDIA GPU with CUDA cores, set ‘use_gpu=Truè and enable PyTorch with the following code block. Otherwise, set ‘use_gpu=Falsè or skip this part.

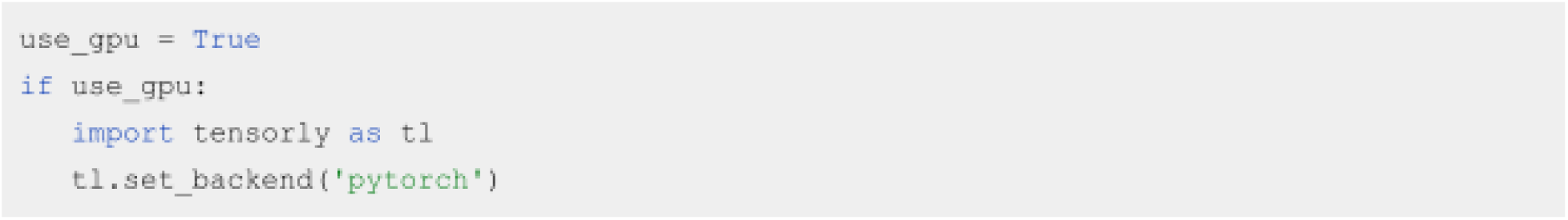

Then, import all the packages we will use in this tutorial:

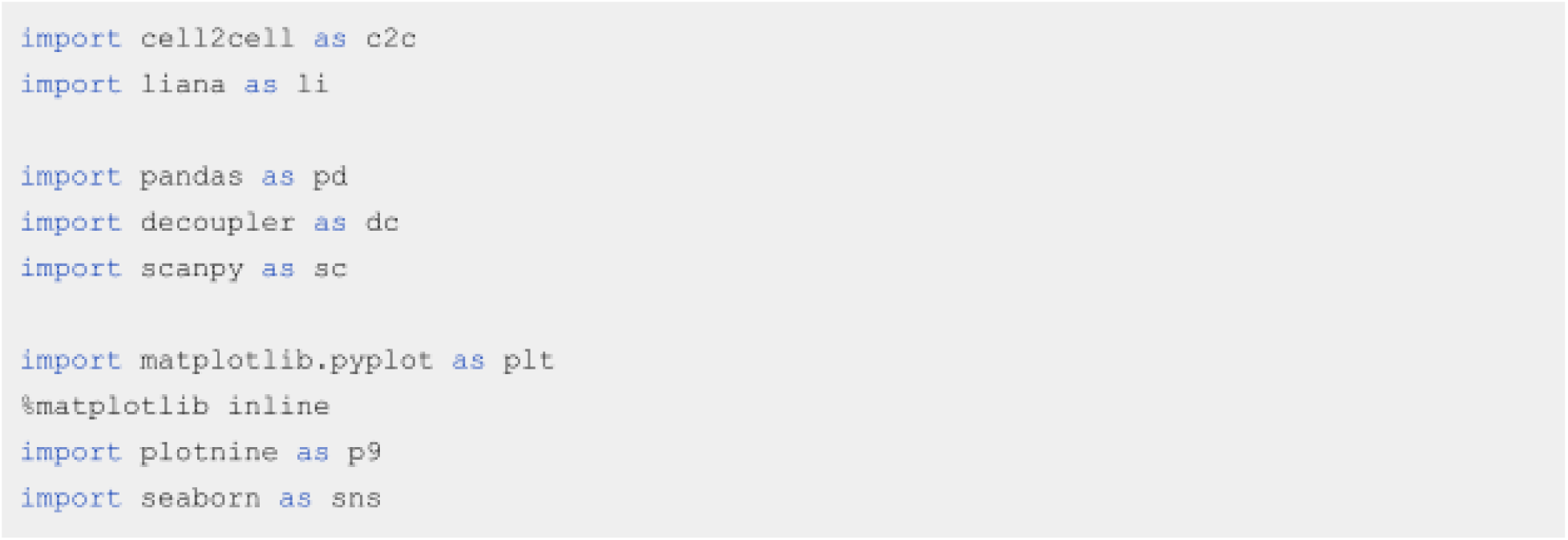

Afterwards, specify the data and output directories:

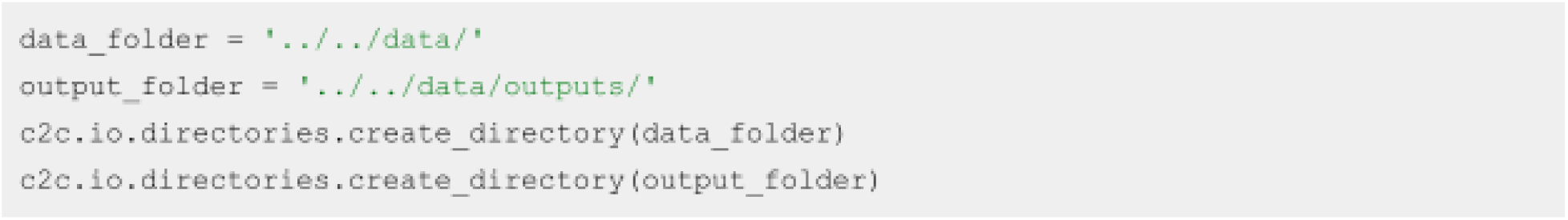

We begin by loading the single-cell transcriptomics data. For this tutorial, we will use a lung dataset of 63k immune and epithelial cells across three control, three moderate, and six severe COVID-19 patients^25^. We use a convenient function to download the data and store it in the AnnData format, on which the scanpy^26^ package is built.

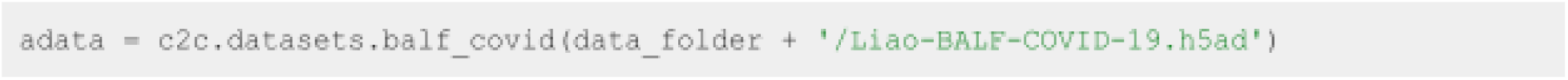

**3. Data Preprocessing**

Data preprocessing is crucial for the correct application of this (**Fig.2a**). Here, we only highlight the essential steps. However, other aspects of data preprocessing should be considered and performed according to the best practices of single-cell analysis (https://github.com/theislab/single-cell-best-practices).

*3.1. Quality Control ● TIMING < 5 min*

The loaded data has already been pre-processed to a degree and comes with cell annotations. Nevertheless, we highlight some of the key steps. To mitigate noise, we filter non-informative cells and genes:

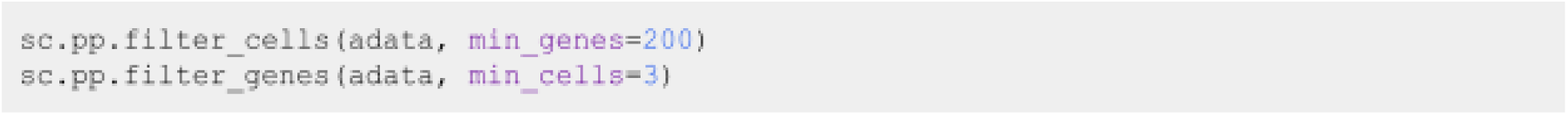

We additionally remove a high mitochondrial content:

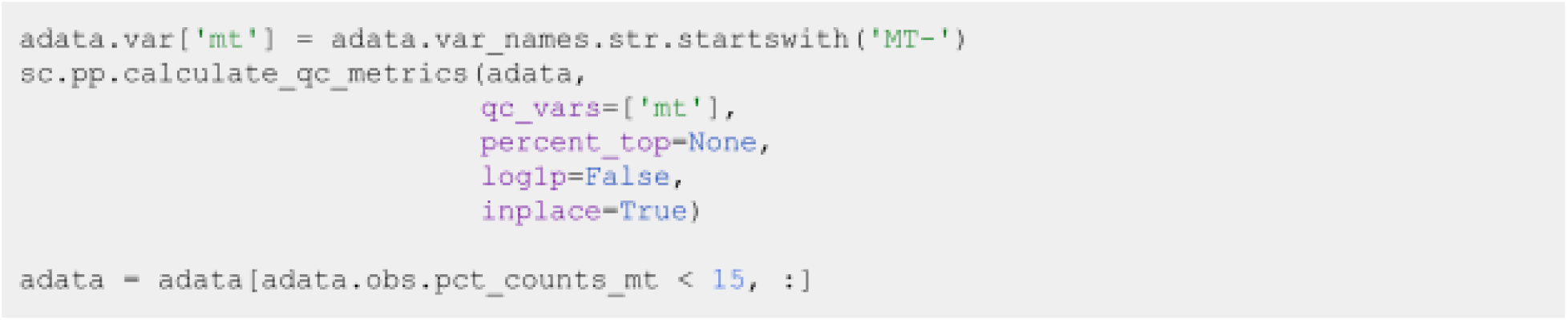

Which is followed by removing cells with a high number of total UMI counts, potentially representing more than one single cell (doublets):

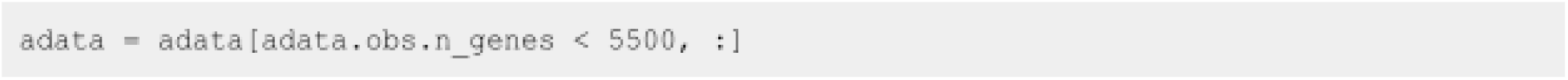

**! CAUTION** Here, we covered the absolute basics. We omit other common practice steps, such as the removal of cells with high ribosomal content and the correction of ambient RNA. Additionally, in certain scenarios, particularly in such where technical variation is expected to be notable, the application of quality control steps by sample is desirable.

*3.2. Normalization ● TIMING < 2 min*

We have now removed the majority of noisy readouts and we can proceed to count normalization, as most cell-cell communication tools typically use normalized count matrices as input. Normalized counts are usually obtained in two essential steps, the first being count depth scaling which ensures that the measured count depths are comparable across cells. This is then usually followed up with log1p transformation, which stabilizes the variance of the counts and enables the use of linear metrics downstream:

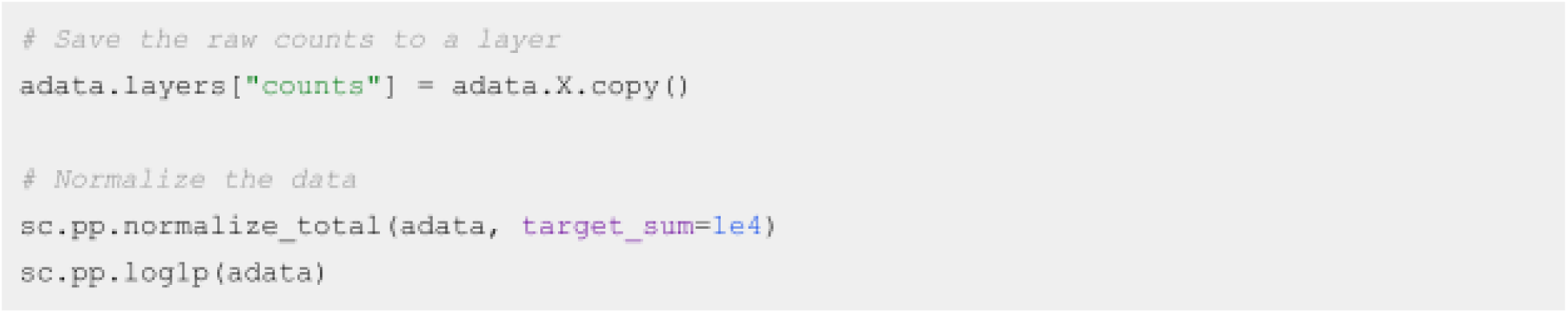

**Δ CRITICAL A key parameter of this command is:**

- **target_sum** ensures that after normalization each observation (cell) has a total count equal to that number.

These normalization steps ensure that the aggregation of cells into cell types, a common practice for CCC inference, is done on comparable cells with approximately normally-distributed feature values.

**? TROUBLESHOOTING** Expression matrices with nan or inf values causes errors. Users should stick to common normalization techniques, and any nan, negative or inf values must be filled to avoid errors.

**4. Inferring cell-cell communication**

Following preprocessing of the single-cell transcriptomics data, we proceed to the inference of potential CCC events (**Fig.3b**). In this case, we will use LIANA to infer the ligand-receptor interactions for each sample. LIANA is available in Python and R, and supports Scanpy, SingleCellExperiment and Seurat objects as input. LIANA is highly modularized, and it natively implements the formulations of several methods, including CellPhoneDBv2^27^, Connectome^28^, log2FC, NATMI^29^, SingleCellSignalR^30^, CellChat^31^, a geometric mean, as well as a consensus score in the form of a rank aggregate^32^ from any combination of methods (**Fig.3**). The high modularity of LIANA further enables the straightforward addition of any other ligand-receptor method.

**Figure 3.**
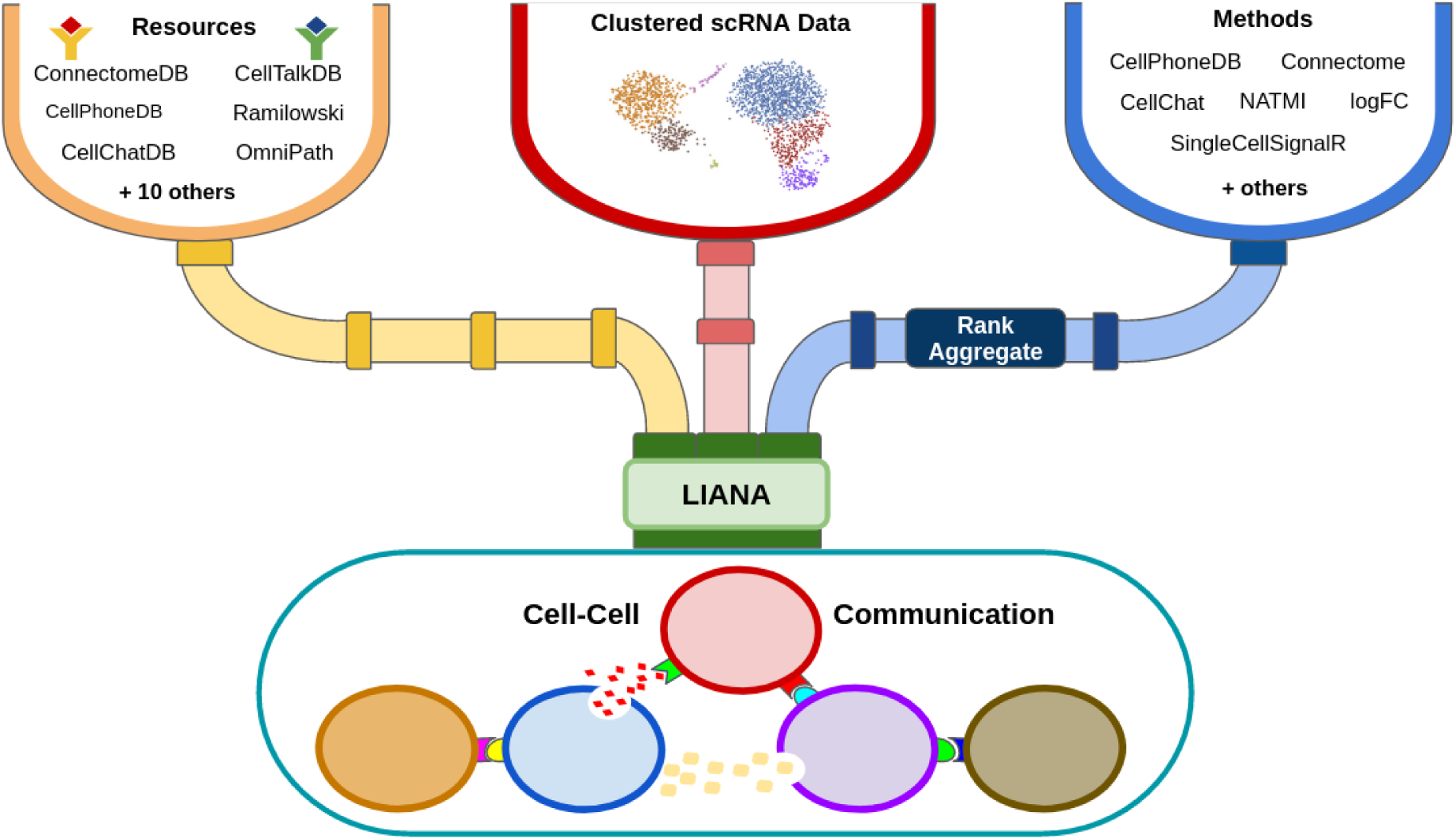
LIANA is a user-friendly and modular ligand-receptor analysis framework. LIANA provides a variety of methods and resources to infer cell-cell communication, making it easy to use multiple existing methods in a coherent manner. It also provides consensus scores and resources to provide generalized results. Figure adapted from^3^.

LIANA classifies the scoring functions from the different methods into two categories: those that infer the “*Magnitude*” and “*Specificity*” of interactions. The “*Magnitude*” of an interaction is a measure of the strength of the interaction, and the “*Specificity*” of an interaction is a measure of how specific an interaction is to a given pair of cell groups. Generally, these categories are complementary, and the magnitude of the interaction is often in agreement with the specificity of the interaction. In other words, a ligand-receptor interaction with a high magnitude score in a given pair of cell types is likely to also be specific, and vice versa.

*4.1. Selecting a method to infer cell-cell communication*

While there are many commonalities between the different methods implemented in LIANA, there also are many variations and different assumptions affecting how the magnitude and specificity scores are calculated (See **Appendix 1**). These variations can result in limited agreement in inferred predictions when using different CCC methods^3, 18, 19^. To this end, in LIANA we additionally provide a rank_aggregate score, that can be used to aggregate any of the scoring functions above into a consensus score.

By default, LIANA calculates an aggregate rank using a re-implementation of the RobustRankAggregate method^32^, and generates a probability distribution for ligand-receptors that are ranked consistently better than expected under a null hypothesis (See **Appendix 1**). The consensus of ligand-receptor interactions across methods can therefore be treated as a P-value. We show in detail how LIANA’s rank aggregate or any of the individual methods can be used to infer communication events from a single sample or context at “Python Tutorial 02 Infer-Communication-Scores” [https://ccc-protocols.readthedocs.io/en/latest/notebooks/ccc_python/02-Infer-Communication-S cores.html].

**Δ CRITICAL** When using LIANA with Tensor-cell2cell, we recommend selecting a scoring function that reflects the *Magnitude* of the interactions, as how the interactions *Specificity* relates to changes across samples is unclear. In this protocol, we will use the ‘*magnitude_rank‘* scoring function from LIANA, under the assumption that ensemble approaches are potentially less biased than any single method alone^20^.

**? TROUBLESHOOTING** The default decomposition method of Tensor-cell2cell is a non-negative Tensor Component Analysis, which, as implied, expects non-negative values as the inputs. Thus, when selecting the method of choice, make sure that you do not have negative CCC scores. If so, you can replace them by zeros or the minimum positive value.

*4.2. Selecting ligand-receptor resources*

When considering ligand-receptor prior knowledge resources, a common theme is the trade-off between coverage and quality, and similarly each resource comes with its own biases^3^. LIANA takes advantage of OmniPath^33^, which includes expert-curated resources of CellPhoneDBv2^27^, CellChat^31^, ICELLNET^34^, connectomeDB2020^29^, CellTalkDB^35^, as well as 10 others^3, 33^. LIANA further provides a consensus expert-curated resource from the aforementioned five resources, along with some curated interactions from SignaLink^36^. In this protocol, we will use the consensus resource from LIANA, though any of the other resources are available via LIANA, and one can also use LIANA with their own custom resource.

Selecting any of the lists of ligand-receptor pairs in LIANA can be done through the following command:

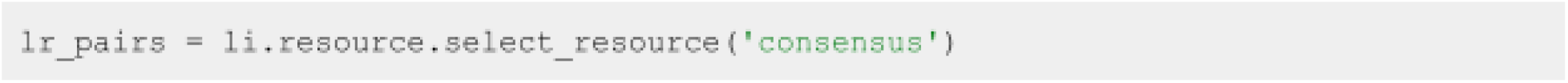

Here ‘consensus’ indicates the use of LIANA’s consensus resource, but it can be replaced by any other available resource (e.g. ‘cellphonedb’, ‘cellchatdb’, ‘connectomeDB’, etc.).

**? TROUBLESHOOTING** Users should choose a resource with gene identifiers and organism that corresponds to that of their data. By default, LIANA uses human gene symbol identifiers, but additionally provides a murine resource as well as functionalities to convert via orthology to other organisms.

*4.3. Running LIANA for each sample ● Timing 4 minutes*

Here, we will run LIANA’s ‘rank_aggregatè with six methods (by default, CellPhoneDBv2, CellChat, SingleCellSignalR, NATMI, Connectome, log2FC) on all of the samples in the dataset.

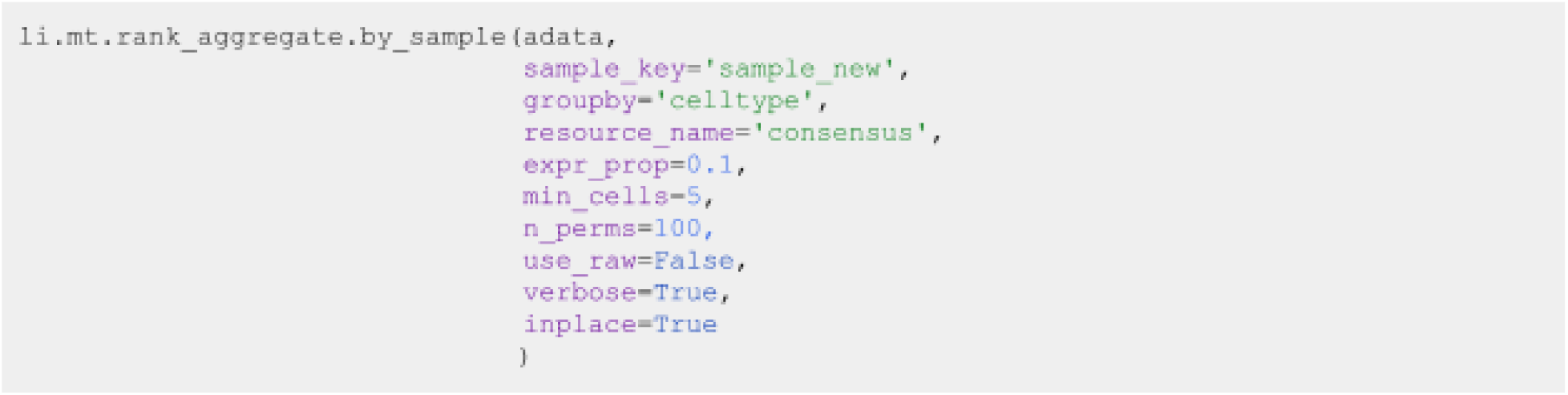

**Δ CRITICAL Key parameters here are:**

- **adata** stands for Anndata, the data format used by scanpy^26^, and we pass here with an object with a single sample/context.
- **sample_key** corresponds to the sample identifiers, available as a column in the ‘adata.obs‘ dataframe.
- **groupby** corresponds to the cell group label stored in ‘adata.obs‘.
- **resource_name** - name of any of the resources available via LIANA
- **expr_prop** is the expression proportion threshold (in terms of cells per cell type expressing the protein) for any protein subunit involved in the interaction, according to which we keep or discard the interactions.
- **min_cells** is the minimum number of cells per cell type required for a cell type to be considered in the analysis
- **n_perms** is the number of permutations for P-value estimation
- **use_raw** is a boolean that indicates whether to use the ‘adata.raw‘ slot, here the log-normalized counts are assigned to ‘adata.X‘, other options include passing the name of a layer via the ‘layer‘ parameter or using the counts stored in ‘adata.raw‘.
- **verbose** is a Boolean value that indicates whether to print the progress of the function
- **inplace** indicates whether storing the results in place, i.e. to ‘adata.uns[“liana_res”]‘.

Δ CRITICAL LIANA considers interactions as occurring only if the ligand and receptor, and all of their subunits, are expressed in at least 10% of the cells (by default) in both clusters involved in the interaction. Any interactions that do not pass these criteria are not returned by default. To return those, the user can use the ‘return_all_lrs‘ parameter. These results will later be used to generate a tensor of ligand-receptor interactions across contexts that will be decomposed into CCC patterns by Tensor-Cell2cell. Thus, how non-expressed interactions are handled is critical to consider when building the tensor later on (see “Python Tutorial 03 Generate-Tensor” [https://ccc-protocols.readthedocs.io/en/latest/notebooks/ccc_python/03-Generate-Tensor.html]).

### Visualize output

One can visualize the output as a dotplot, but with the addition of the samples.

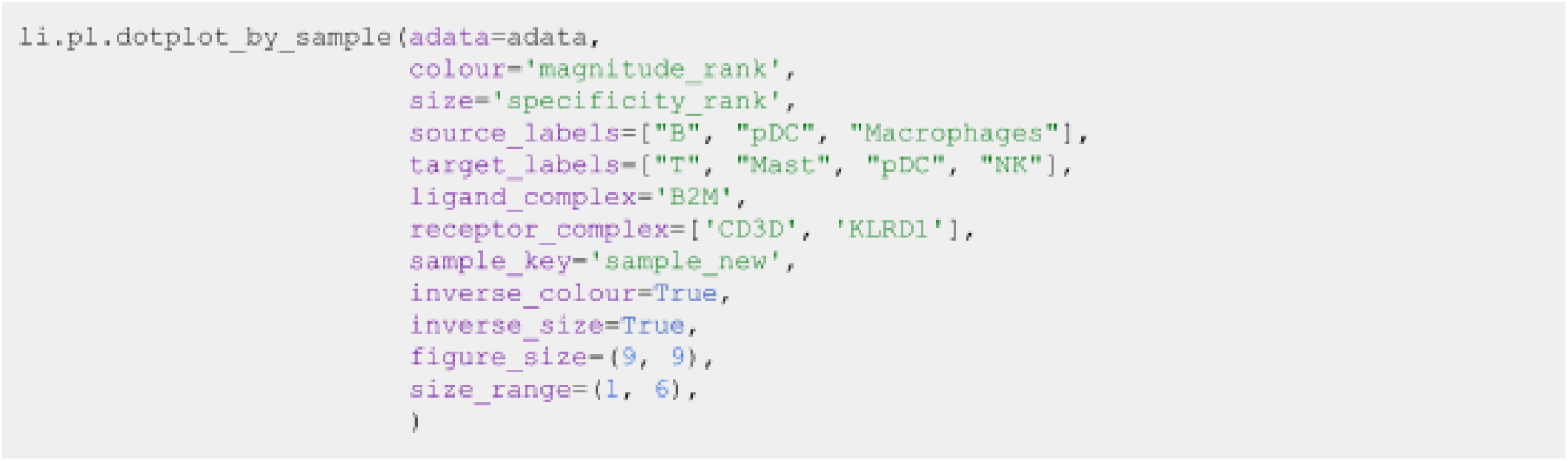

**Key parameters here are:**

- **source_labels** is a list containing the names of the sender cells of interest.
- **target_labels** is a list containing the names of the receiver cells of interest.
- **ligand_complex** is a list containing the names of the ligands of interest.
- **receptor_complex** is a list containing the names of the receptors of interest.
- **sample_key** is a string containing the column name where samples are specified.

**▪ PAUSE POINT** We can export the liana results by sample to a csv, and save them for later use:

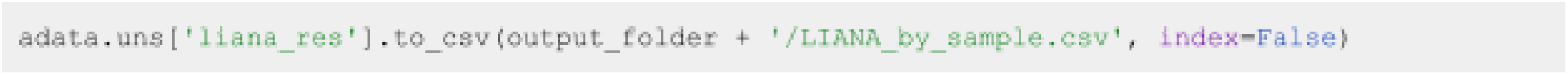

Alternatively one could just export the whole AnnData object, together with the ligand-receptor results stored at ‘adata.uns[‘liana_res’]‘:

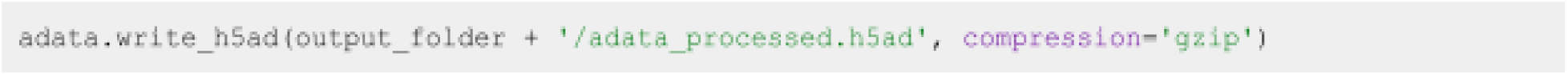

**5. Comparing cell-cell communication across multiple samples**

*5.1. Building a 4D-communication tensor ● Timing <1 minute*

First, we generate a list containing all samples from our AnnData object. For visualization purposes we sort them according to COVID-19 severity. Here, we can find the names of each of the samples in the ‘sample_new’ column of the adata.obs information:

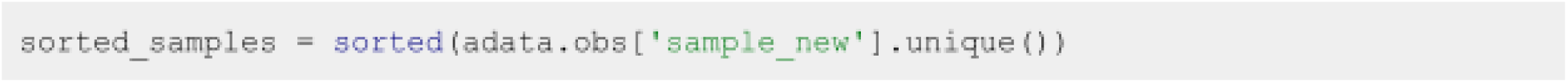

Tensor-cell2cell performs a tensor decomposition to find context-dependent patterns of cell-cell communication. It builds a 4D-communication tensor, which in this case is built from the communication scores obtained from LIANA for every combination of ligand-receptor and sender-receiver cell pairs per sample (**Fig.2c**). For this protocol and associated tutorials, we implemented a function that facilitates building this communication tensor:

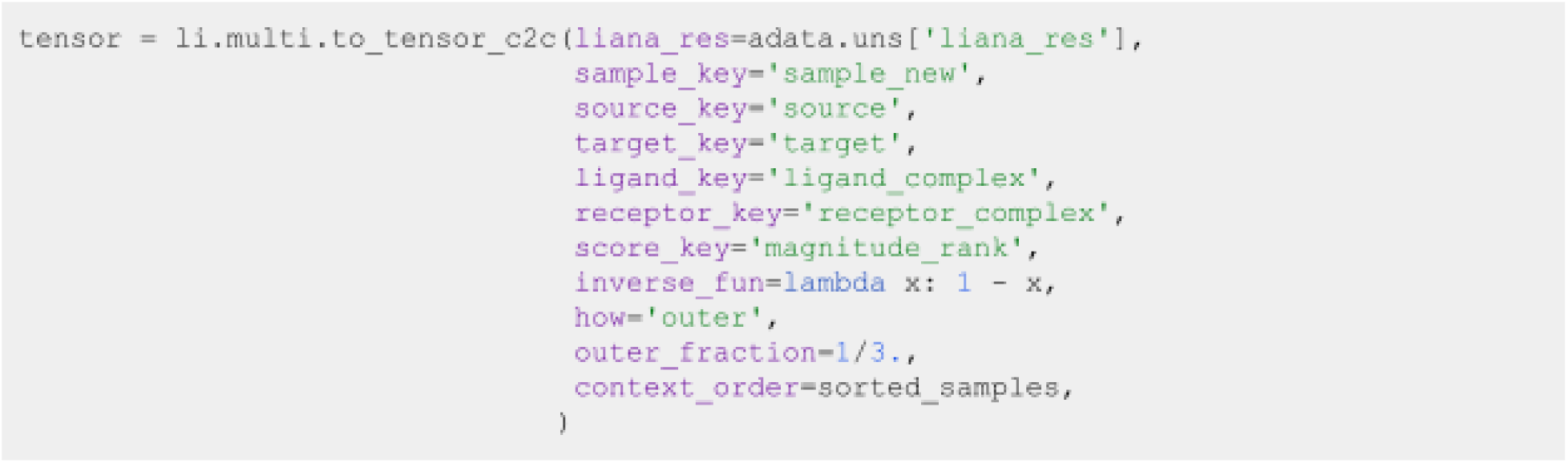

**? TROUBLESHOOTING** Since the ‘*magnitude_rank*‘ from LIANA represents a score where the values closest to 0 represent the most probable communication events, we need to invert the communication scores to use it with Tensor-cell2cell. See the parameter ‘inverse_fun‘ below for further details for transforming this score.

**Δ CRITICAL Key parameters here are:**

- **liana_res** is the dataframe containing the results from LIANA, usually located in ‘adata.uns[‘liana_res’]. We can pass directly the AnnData object to the parameter adata to this function. If the AnnData object is passed, we do not need to specify the liana_res parameter.
- **sample_key**, **source_key**, **target_key**, **ligand_key**, **receptor_key**, and **score_key** are the column names in the dataframe containing the samples, sender cells, receiver cells, ligands, receptors, and communication scores, respectively. Each row of the dataframe contains a unique combination of these elements.
- **inverse_fun** is the function we use to convert the communication score before building the tensor. In this case, the ’magnitude_rank’ score generated by LIANA considers low values as the most important ones, ranging from 0 to 1. In contrast, Tensor-cell2cell requires higher values to be the most important scores, so here we pass a function (lambda x: 1 - x) to adapt LIANA’s magnitude-rank scores (subtracts the LIANA’s score from 1). If None is passed instead, no transformation will be performed on the communication score. If using other scores coming from one of the methods implemented in LIANA, a similar transformation can be done depending on the parameters and assumptions of the scoring method.
- **how** controls which ligand-receptor pairs and cell types to include when building the tensor. This decision depends on whether the missing values across a number of samples for both ligand-receptor interactions and sender-receiver cell pairs are considered to be biologically-relevant. Options are:
  `o`***’inner’*** is the most strict option since it only considers cell types and ligand-receptor pairs that are present in all contexts (intersection).
  `o`***’outer’*** considers all cell types and ligand-receptor pairs that are present across contexts (union).
  `o`***’outer_lrs’*** considers only cell types that are present in all contexts (intersection), while all ligand-receptor pairs that are present across contexts (union).
  `o`***’outer_cells’*** considers only ligand-receptor pairs that are present in all contexts (intersection), while all cell types that are present across contexts (union).
- **outer_fraction** controls the elements to include in the union scenario of the how options. Only elements that are present at least in this fraction of samples/contexts will be included. When this value is 0, the tensor includes all elements across the samples. When this value is 1, it acts as using how=’inner’.
- **context_order** is a list specifying the order of the samples. The order of samples does not affect the results, but it is useful for posterior visualizations.

We can check the shape of this tensor to verify the number of samples, ligand-receptor pairs, sender cells, and receiver cells, respectively:

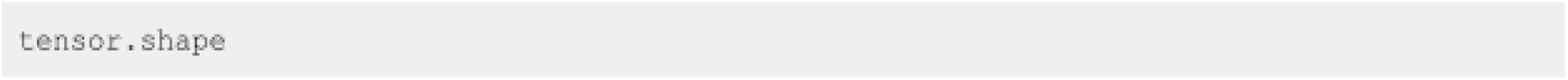

In addition, optionally we can generate the metadata for coloring the elements in each of the tensor dimensions (i.e., for each of the contexts/samples, ligand-receptor pairs, sender cells, and receiver cells) in posterior visualizations. This metadata corresponds to dictionaries for each of the dimensions, containing the elements and their respective major groups, such as a signaling categories for a ligand-receptor interactions, a hierarchically more granular cell type, or a disease condition for a sample. In cases where we do not account for such information, we do not need to generate such dictionaries.

For example, we can build a dictionary for the contexts/samples dictionary by using the metadata in the AnnData object. In this example dataset, we can find samples in the column ‘sample_new’, while their majors groups (representing COVID-19 severity) are found in the column ‘condition’:

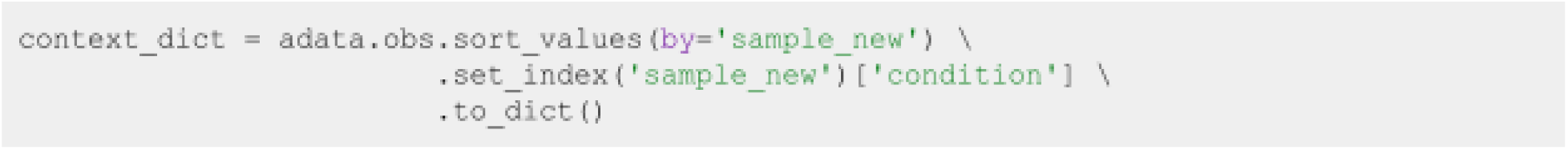

Then, the metadata can be generated with:

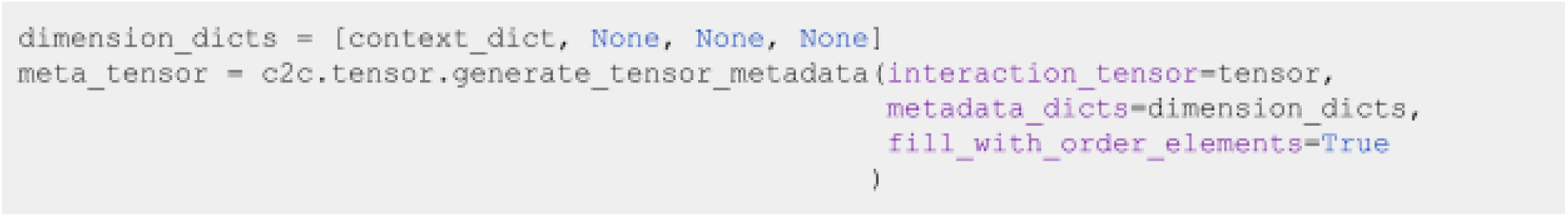

Notice that the None elements in the variable dimensions_dicts represent the dimensions where we are not including additional metadata. If you want to include metadata about major groups for those dimensions, you have to replace the corresponding None by a dictionary as described before.

**▪ PAUSE POINT** We can export our tensor and its metadata for performing the tensor decomposition later:

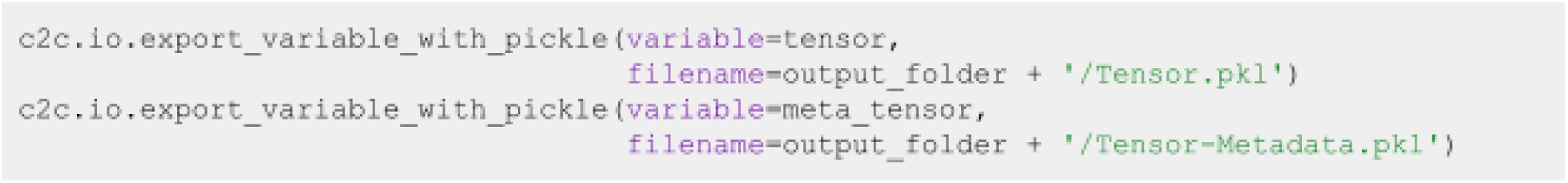

Then, we can load them with:

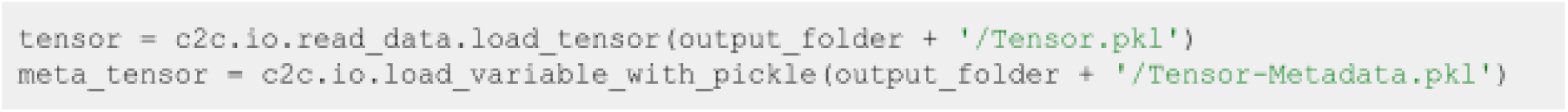

*5.2. Running Tensor-cell2cell across samples ● Timing 5 minutes with a ‘regular’ run or 40 minutes with a ‘robust’ run (using a GPU in both cases)*

Now that we have built the tensor and its metadata, we can run Tensor Component Analysis via Tensor-cell2cell with one simple command that we implemented for our unified tools:

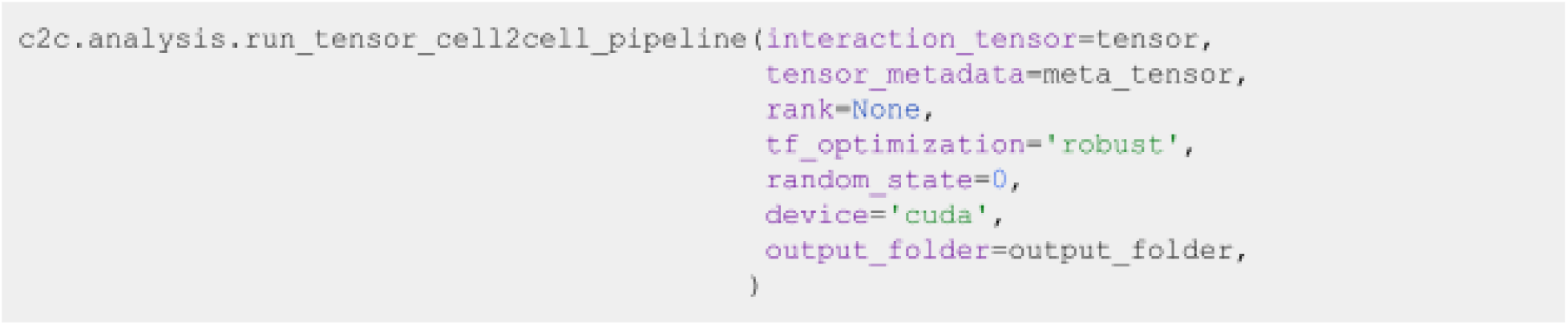

**Δ CRITICAL Key parameters of this command are:**

- **rank** is the number of factors or latent patterns we want to obtain from the analysis. You can either indicate a specific number or leave it as None to perform the decomposition with a suggested number from an elbow analysis.
- **tf_optimization** indicates whether running the analysis in the ’regular’ or the ’robust’ way. The regular way runs the tensor decomposition fewer times than the robust way to select an optimal result. Additionally, the former employs less strict convergence parameters to obtain optimal results than the latter, which is also translated into a faster generation of results.
- **random_state** is the seed for randomization. It controls the randomization used when initializing the optimization algorithm that performs the tensor decomposition. It is useful for reproducing the same result every time that the analysis is run. If None, a different randomization will be used each time.
- **device** indicates whether we are using the ’cpu’ or a GPU with ’cuda’ cores. See the Installation section of this tutorial for instructions to enable the use of GPU(s).
- **output_folder** is the full path to the folder where the results will be saved. Make sure that this folder exists before passing it here.

This command will output three main results: a figure with the elbow analysis for suggesting a number of factors (only if rank=None), a figure with the loadings assigned to each element

within a tensor dimension per factor obtained, and an excel file containing the values of these loadings.

**? TROUBLESHOOTING** Elbow analysis returns a rank equal to one, or the curve is increasing instead of decreasing. This may be due to high sparsity in the tensor. The sparsity can be decreased by re-building the 4D tensor after re-running LIANA (**Step 4.3**) with a smaller ‘expr_prop‘ (e.g. ‘expr_prop=0.05‘) or by only re-building the tensor (**Step 5.1**) with a higher ‘outer_fraction‘ (e.g. ‘outer_fraction=0.8‘).

*5.3. Downstream visualizations: Making sense of the factors ● Timing <2 minutes*

The figure representing the loadings in each factor generated in the previous section can be interpreted by interconnecting all dimensions within a single factor. For example, if we take Factor 4 in **Fig.4**, the CCC program here occurs in each sample in a manner proportional to their loadings, here correlated with COVID-19 severity. Relevant interactors can be interpreted according to their loadings (i.e. ligand-receptor pairs, sender cells, and receiver cells with high loadings play a more prominent role in an identified CCC program). Ligands in high-loading ligand-receptor pairs are sent predominantly by high-loading sender cells, and interact with the cognate receptors on the high-loadings receiver cells. In this factor, the program would be predominantly driven by changes in the receptor expression of receiver cells such as macrophages, neutrophils and myeloid dendritic cells.

**Figure 4.**
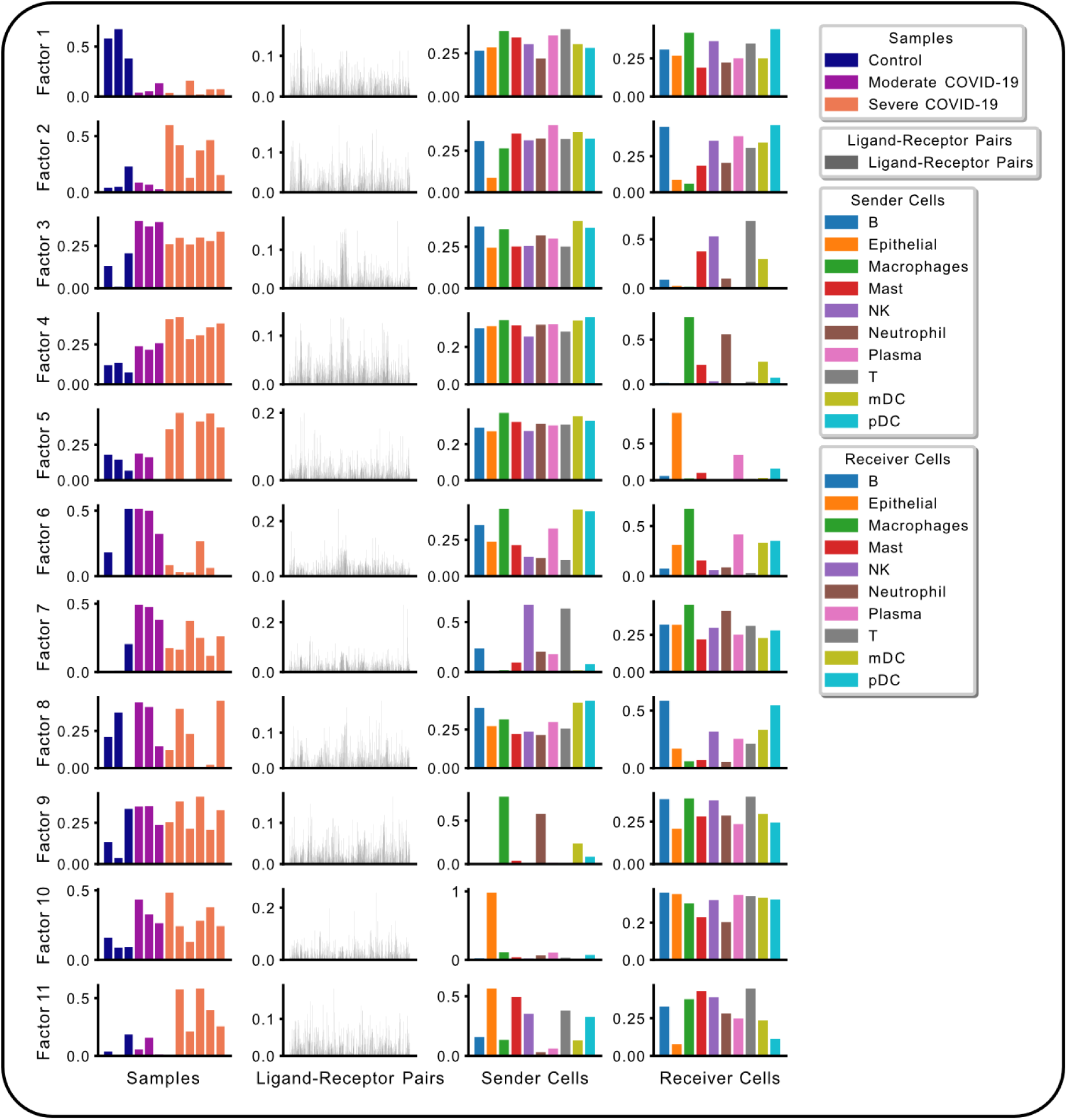
Cell-cell communication programs obtained by combining LIANA and Tensor-cell2cell. After inferring cell-cell communication with LIANA from the COVID-19 data, and running a Tensor Component Analysis with Tensor-cell2cell, 11 factors were obtained (rows here), each of which represents a different cell-cell communication program. Within a factor, loadings (y-axis) are reported for each element (x-axis) in every tensor dimension (columns). Elements here are colored by their major groups as indicated in the legend.

We can access the loading values of samples in each of the factors with:

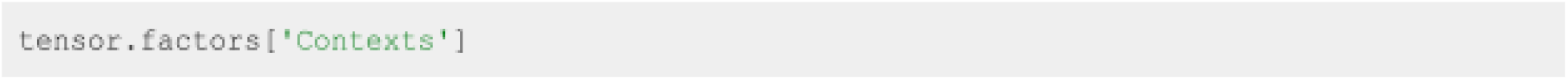

In this case we obtain a dataframe where rows represent the samples, columns the factors generated by the decomposition, and entries are the loadings of each element within the corresponding factor. We can also access the loadings for the elements in the other dimensions by replacing ‘Contexts’ with ‘Ligand-Receptor Pairs’, ‘Sender Cells’, or ‘Receiver Cells’. Then, we can use these loadings to perform various downstream analyses (**Fig.5**).

**Figure 5.**
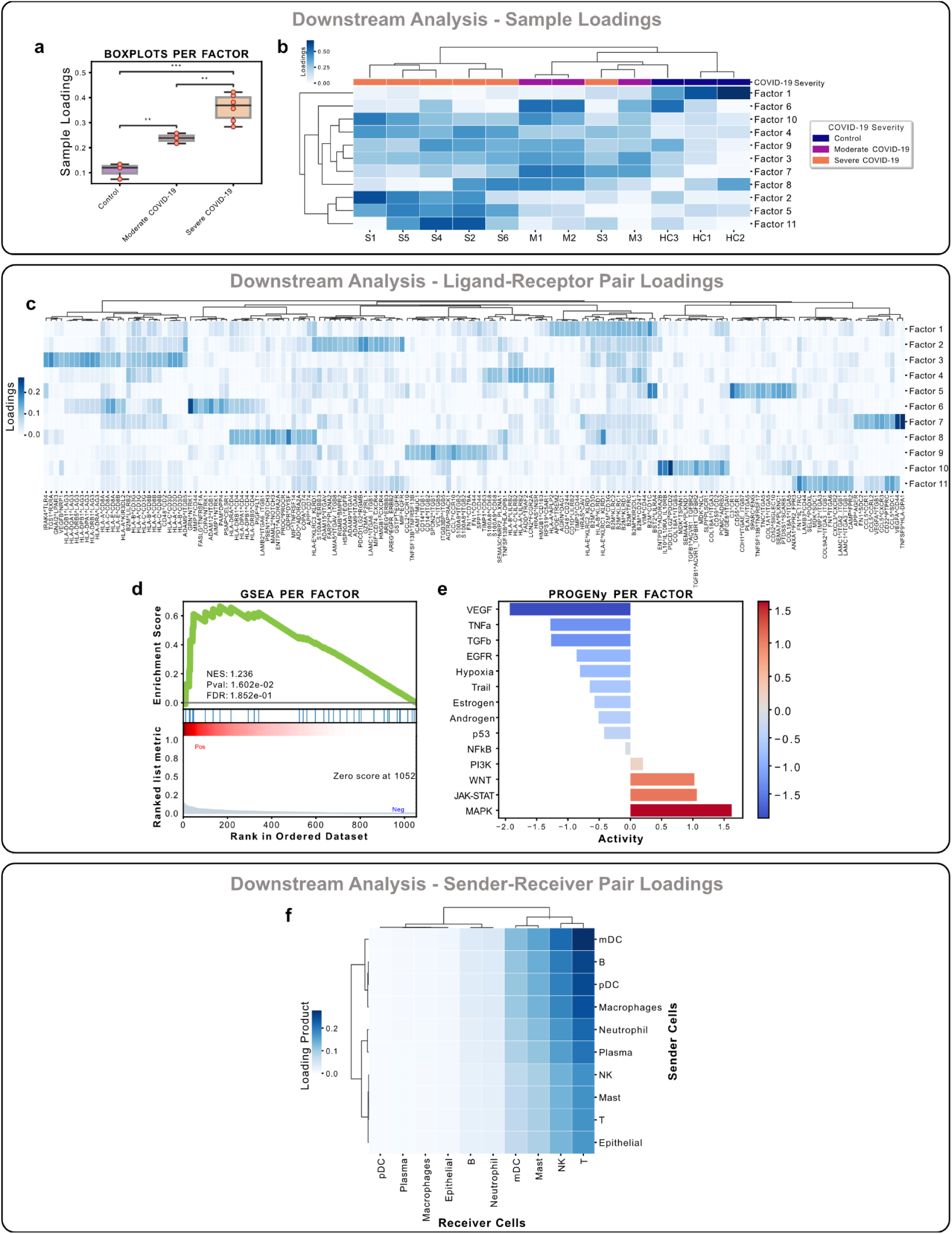
Examples of downstream analyses performed on the results from the LIANA and Tensor-cell2cell framework. Downstream analyses can be performed by using the loadings of one of the tensor dimensions. Context or sample loadings (**a-b**) can be used to (**a**) compare statistically different condition groups within the same cell-cell communication program or (**b**) to group samples across all programs. (**c-e**) Similarly, ligand-receptor interactions can be analyzed from their loadings per or across factors. (**c**) Key ligand-receptor pairs whose loadings are above a threshold can be clustered depending on their importance across all cell-cell communication programs. They can also be ranked according to their loadings within a factor (factor-specific analyses), and this information can be used to run an enrichment analysis such as (**d**) GSEA or (**e**) PROGENy to associate each of the programs with different functions or pathways. (**f**) Finally, cell type loadings can be jointly used within a factor to have an overall representation of the cell-cell communication (i.e., a factor-specific network of communication).

For example, we can use loadings to compare groups of samples (**Fig.5a-b**) with box plots and statistical tests:

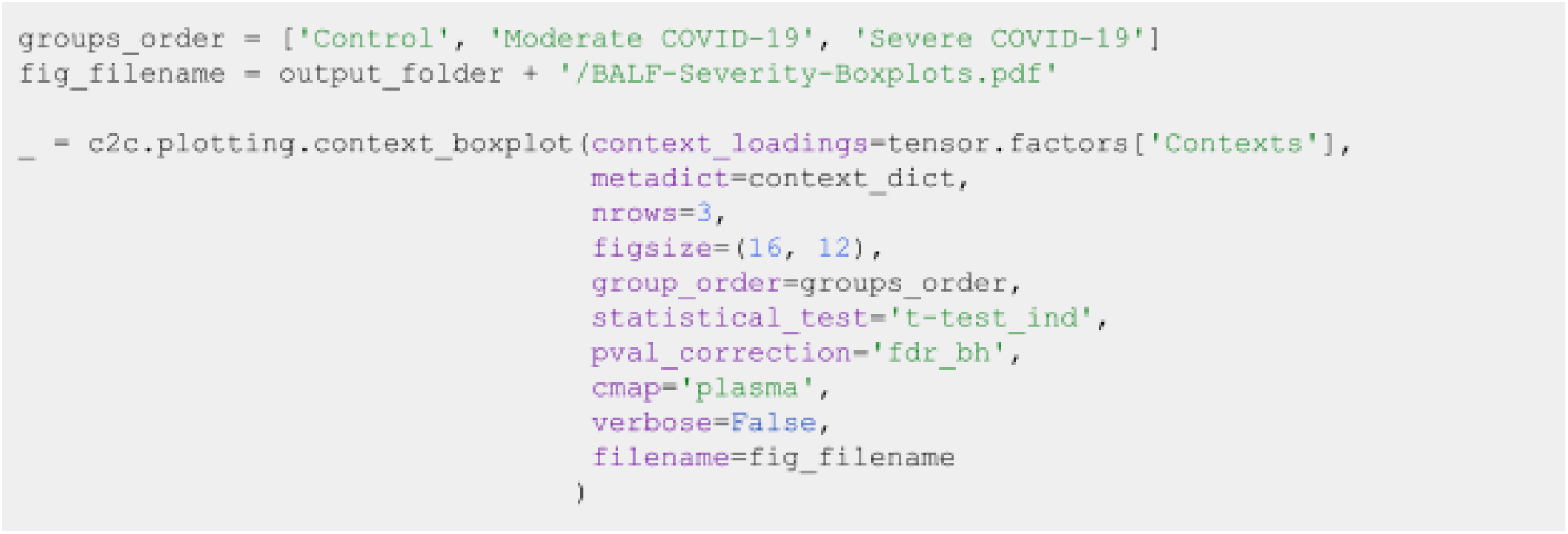

**Δ CRITICAL** In this case, we can change the statistical test and the multiple-test correction with the parameters ‘statistical_test‘ and ‘pval_correction‘. Here we used an independent t-test and a Benjamini-Hochberg correction. Additionally, we can set verbose=True to print exact test statistics and P-values.

We can also generate heatmaps for the elements with loadings above a certain threshold in a given dimension (**Fig.5b,c,f**). Furthermore, we can cluster these elements by the similarity of their loadings across all factors:

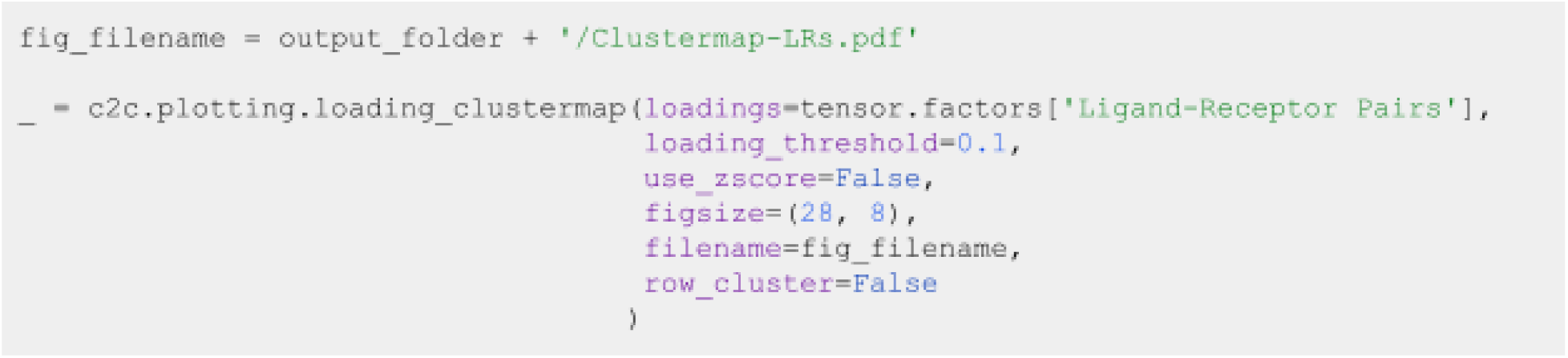

**? TROUBLESHOOTING** Note that here we plot the loadings of the dimension representing the ligand-receptor pairs. In addition, we prioritize the pairs with high loadings using the parameter ‘loading_threshold=0.1‘. In this case, the elements are included only if they are greater than or equal to that threshold in at least one of the factors. If we use ‘loading_threshold=0‘, we would consider all of the elements. Considering all of the elements would require modifying the parameter ‘figsizè to enlarge the figure.

**! CAUTION** Changing the parameter ‘use_zscorè to **True** would standardize the loadings of one element across all factors. This is useful to compare an element across factors and highlight the factors in which that element is most important. Modifying ‘row_cluster‘ to **True** would also cluster the factors depending on the elements that are important in each of them.

**6. Pathway Enrichment Analysis: Interpreting the context-driven communication**

The decomposition of ligand-receptor interactions across samples into loadings associated with the conditions reduces the dimensionality of the inferred interactions substantially. Nevertheless, we are still working with 1,054 interactions across multiple factors associated with the disease labels. To this end, as is commonly done when working with omics data types, we can perform pathway enrichment analysis to identify the general biological processes of interest. By using the loadings for each ligand-receptor pair, we can rank them within each factor and use this ranking as input to enrichment analysis (**Fig.5d-e**). Pathway enrichment thus serves two purposes; it further reduces the dimensionality of the inferred interactions, and it enhances the biological interpretability of the inferred interactions.

Here, we will show the application of classical gene set enrichment analysis on the ligand-receptor loadings. We will use GSEA^37^ with KEGG Pathways^38^, as well as a multivariate linear regression from decoupler-py^39^ with the PROGENy pathway resource^40^.

First, we assign ligand-receptor loadings to a variable:

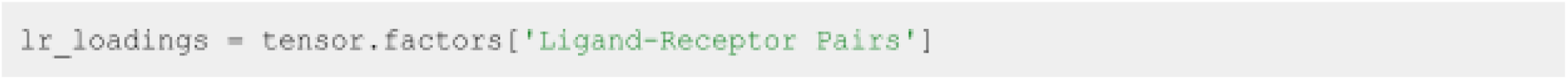

*6.1. Classic Pathway Enrichment*

For the pathway enrichment analysis, we use ligand-receptor pairs instead of individual genes. KEGG was initially designed to work with sets of genes, so first we need to generate ligand-receptor sets for each of its pathways. A ligand-receptor pair is assigned as part of a pathway set if all of the genes in the pair are part of the gene set of such pathway:

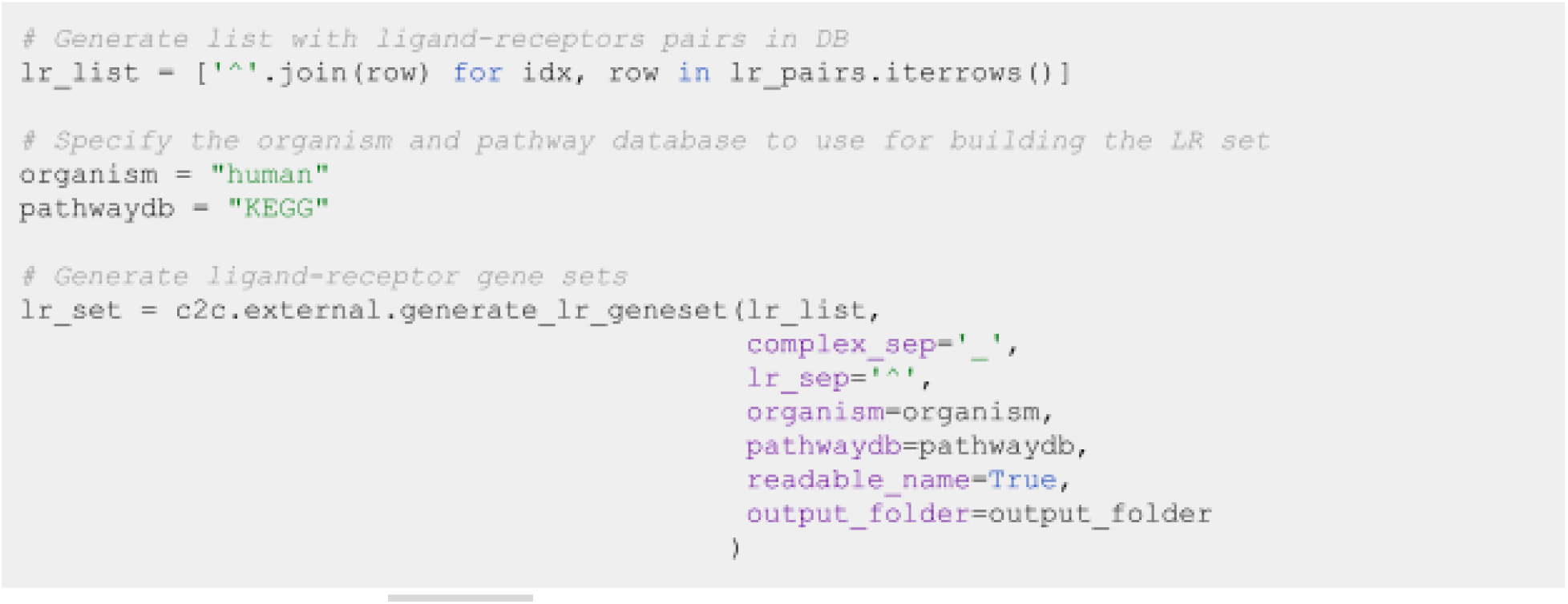

Note that we use the ‘lr_pairs‘ database that we loaded in the ***Selecting ligand-receptor resources*** section.

**Δ CRITICAL Key parameters of this command are:**

- **complex_sep** indicates the symbol separating the gene names in the protein complex.
- **lr_sep** is the symbol separating a ligand and a receptor complex.
- **organism** is the organism matching the gene names in the single-cell dataset. It could be either “human” or “mouse”.
- **pathwaydb** is the name of the database to be loaded, provided with the cell2cell package. Options are “GOBP”, “KEGG”, and “Reactome”.

Run GSEA via cell2cell which calls the ‘gseapy.prerank‘ function internally ● **Timing** < 1 minute

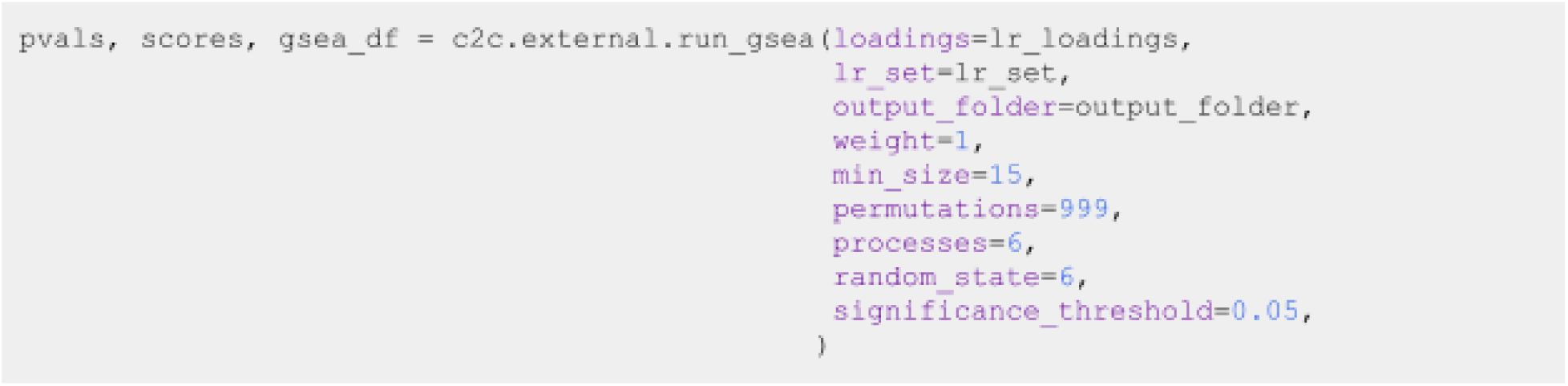

**Δ CRITICAL Key parameters of this command are:**

- **lr_set** is a dictionary associating pathways (keys) with ligand-receptor pairs (values).
- **weight** represents the original parameter p in GSEA. It is an exponent that controls the importance of the ranking values (loadings in our case).
- **min_size** indicates the minimum number of LR pairs that a set has to contain to be considered in the analysis.
- **permutations** indicates the number of permutations to perform to generate the null distribution.
- **random_state** is the reproducibility seed.
- **significance_threshold** is the P-value threshold to consider significance.

Now that we have obtained the normalized-enrichment scores (NES) and corresponding P-values from GSEA, we can plot those using the following function from cell2cell:

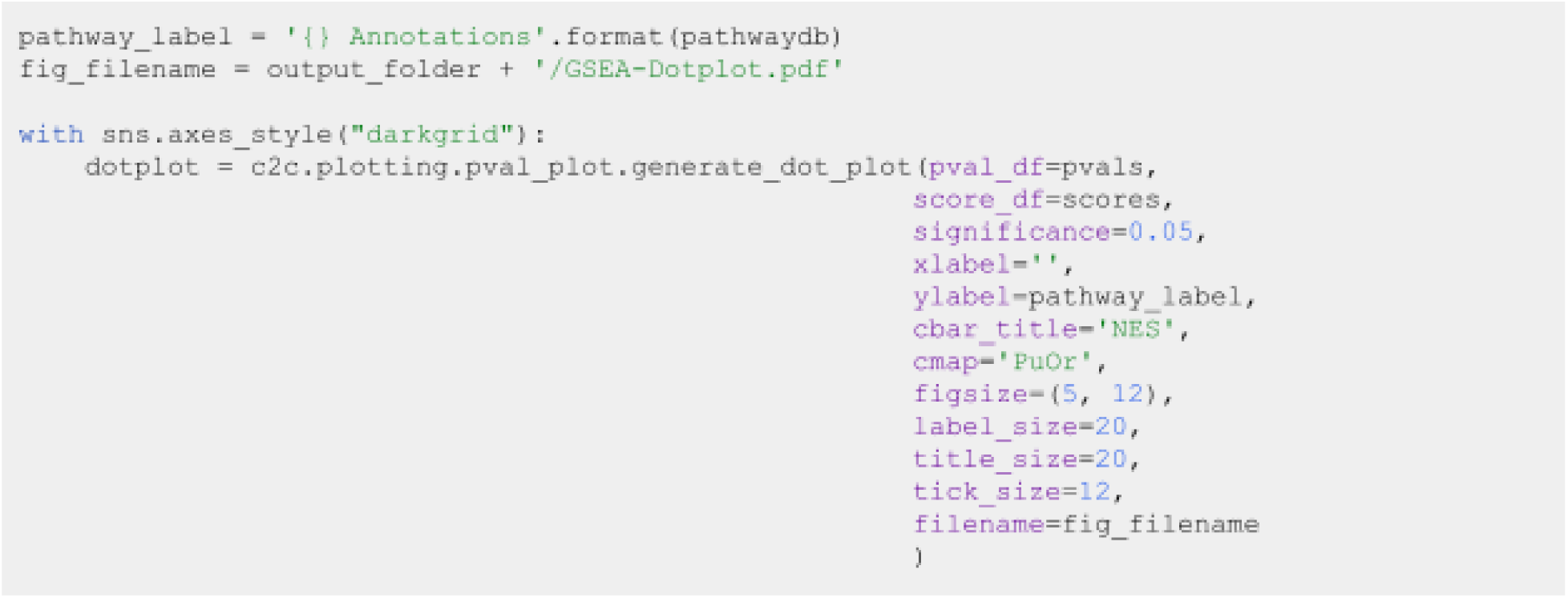

*6.2. Footprint enrichment analysis*

In footprint enrichment analysis, instead of considering the genes whose products (proteins) are directly involved in a process of interest, we consider the genes affected by it - i.e. those that change downstream as a consequence of the process^41^. In this case, we will use the PROGENy resource to infer the pathways driving the identified context-dependent patterns of ligand-receptor pairs. PROGENy was built in a data-driven manner using perturbation data^40^. Consequently, it assigns different weights to each gene in its pathway genesets according to its importance. Thus, we need an enrichment method that can account for weights. To do so, we will use a multivariate linear regression implemented in decoupler-py^39^.

As we did in GSEA using Tensor-cell2cell, we first have to generate ligand-receptor gene sets while also assigning a weight to each ligand-receptor interaction. This is done by taking the mean between the ligand and receptor weights. For ligand and receptor complexes, we first take the mean weight for all subunits. We keep ligand-receptor weights only if all the proteins in the interaction are sign-coherent and present for a given pathway.

Load the PROGENy genesets and then convert them to sets of weighted ligand-receptor pairs:

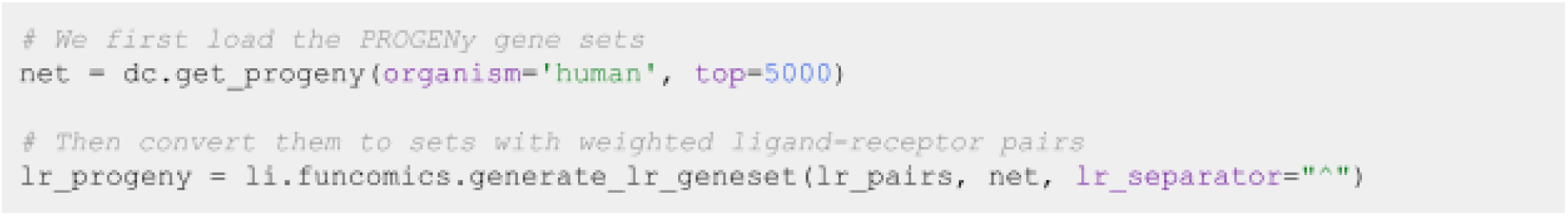

Run footprint enrichment analysis using the ‘mlm‘ method from decoupler-py ● Timing < 1 minute:

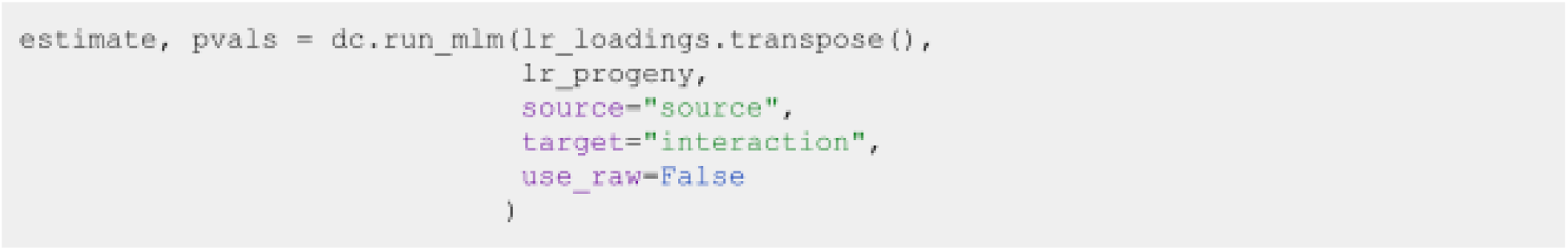

Here, ‘estimatè and ‘pvals‘ correspond to the t-values and P-values assigned to each pathway. Finally, we generate Heatmap for the 14 Pathways in PROGENy across all Factors:

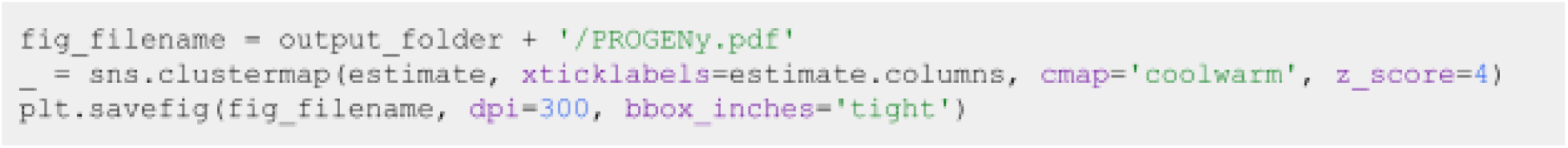

From the heatmap, we can also generate a Barplot for the PROGENy pathways for a specific factor:

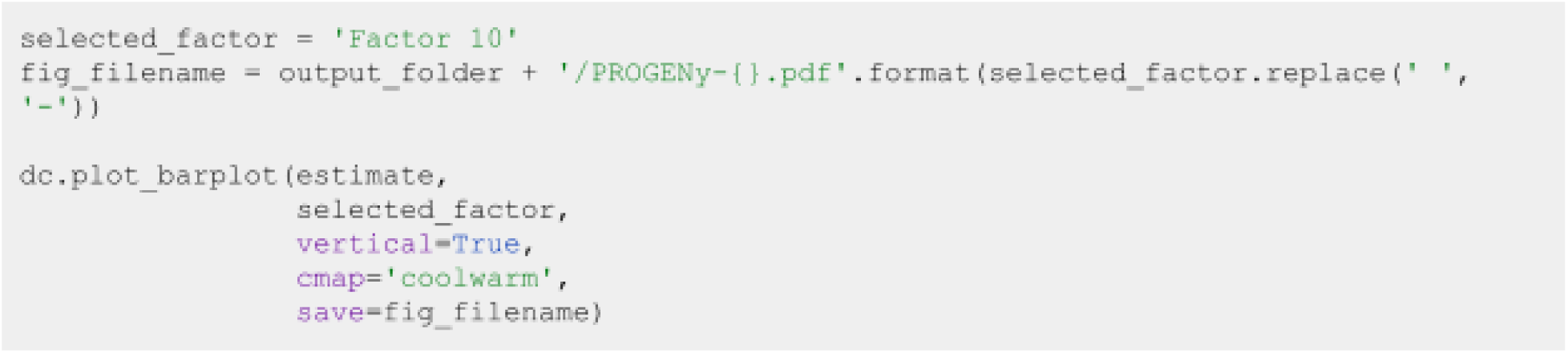

## Troubleshooting

Troubleshooting advice is summarized in Table 2.

**Table 2.** Troubleshooting.

| Step | Problem | Possible reason | Solution |
| --- | --- | --- | --- |
| 3 & 4 | Error: Expression matrix contains non-finite values (nan or inf)<br>Warning: Make sure that normalized counts are passed | Mishandling counts processing | Ensure that the matrix containing normalized counts is passed. Replace nan and inf values by zeros. |
| 4.1 | Negative values in LIANA outputs | Using preprocessed data with negative expression values. | Avoid using preprocessing methods that generate negative values (e.g. centering the data to the mean values, using batch-corrected expression values, etc.). |
| 4.2 | Not enough ligand-receptor pairs in the data for the analysis | Mismatched symbol IDs | LIANA by default uses a resource with gene symbol IDs. When working with e.g. Ensembl IDs users need to provide an external resource; see https://ccc-protocols.readthedocs.io/en/latest/notebooks/ccc_python/02-Infer-Communication-Scores.html |
| 5.1 | CCC scores representing opposed importance | When using 'magnitude_rank' scores from LIANA, lower values are more important. However, Tensor-cell2cell prioritizes high values as the important ones. | Build the 4D tensor using an 'inverse_fun' to make lower values to be the most important scores. |
| 5.2 | Rank selection through the elbow analysis is not behaving properly | High sparsity or number of missing values in the tensor | Re-run LIANA with less stringent parameters (e.g. smaller expr_pror). Re-build the tensor with more strict how parameters (e.g. using how='inner' or increasing outer_fraction). |
| 5.3 | Visualization of loadings are not properly displayed in heatmaps | Too many or few elements in the dimension to visualize | To visualize all elements, use the parameter 'loading_threshold=0' to create the heatmaps. If you have too many elements, you can prioritize those with high loadings, so a threshold can be set. E.g., 'loading_threshold=0.1' |

## Timing

Step 1. Installation of Anaconda/Miniconda and Python packages: 5-30 mins. Step 2. Initial setups: ∼1 min.

Step 3. Deciphering cell-cell communication with LIANA: ∼5 min.

Step 4. Comparing cell-cell communication across multiple samples with Tensor-cell2cell: Rank estimation with elbow analysis takes 5 min, while the tensor decomposition 40 min.

Step 5. Downstream visualizations: 1 min.

Step 6. Functional Enrichment Analysis of KEGG and PROGENy pathways respectively using GSEA and linear regression take 1 min each.

## Anticipated Results

Deciphering cell-cell communication with LIANA yields all ligand-receptor interactions, defined in the prior knowledge resource, for every pair of cell types within the dataset. For each interaction, a set of statistics is assigned. These typically include a value that reflects the magnitude and specificity of interaction depending on the method of choice. The magnitude scores for each interaction in each sample are transformed into a 4D tensor that is then decomposed by Tensor-cell2cell. Prior to decomposition, it is recommended to estimate the optimal number of factors required to reconstruct the original tensor. For each output factor, we obtain four vectors that represent the sample, ligand-receptor interaction, sender cell type, and receiver cell type loadings. We can interpret the loadings as the relative importance of each element in each dimension of the original tensor. Together, the four vectors in a given factor constitute the CCC programs. The vectors are interconnected such that their combination across dimensions define a CCC program, with loadings in the sample dimension representing the context-dependence of the program and elements from each of the other dimensions (ligand-receptor interactions and cell types) with high loadings being key mediators of this program. By focusing on sample loadings associated with a given condition label, we can thus identify the cell types and interactions also associated with that label. To aid the interpretation of LIANA and Tensor-cell2cell results, we also provide a wide range of visualizations and strategies to summarize the interaction loadings into biologically-meaningful insights. We anticipate that our unified protocol will aid the scientific community in studying CCC using large single-cell datasets with a high number of samples and biological conditions.

## Acknowledgements

D.D. is supported by the European Union’s Horizon 2020 research and innovation program (860329 Marie-Curie ITN “STRATEGY-CKD”). E.A. is supported by the Chilean Agencia Nacional de Investigación y Desarrollo (ANID) through its scholarship program DOCTORADO BECAS CHILE/2018 -72190270, the Fulbright Chile Commission, and the Siebel Scholars Foundation. This work was further supported by the NVIDIA Academic Hardware Grant Program. N.E.L. is supported in part by NIGMS R35 GM119850. H.B. is also supported by an ORISE fellowship.

## Conflict of interests

JSR reports funding from GSK, Pfizer and Sanofi and fees from Travere Therapeutics, and Astex. During the course of this work, NEL reports funding from Sanofi, Amgen, Sartorius, and Ionis, and is a co-founder of NeuImmune Inc. and Augment Biologics.

## Authors contributions

H.B., D.D, and E.A. conceived the project, adapted the computational tools, developed the protocol, and wrote the initial version of the manuscript. J.S.R. and N.E.L. revised the manuscript and supervised the project. H.B., D.D, and E.A. contributed equally. J.S.R. and N.E.L are both corresponding authors and have contributed equally.

## Notes

https://github.com/saezlab/ccc_protocols

https://ccc-protocols.readthedocs.io/en/latest/index.html

https://zenodo.org/record/7706962/

