## Supplementary Information for "Combining LIANA and Tensor-cell2cell to decipher cell-cell communication across multiple samples"

### Appendix 1.

With choice comes responsibility, so to guide the selection of a ligand-receptor method, let's have a more detailed look at the scoring functions from the diverse methods implemented in LIANA.

#### Shared Notations

$k$ is the k-th ligand-receptor interaction

$L$ - expression of ligand L

$R$ - expression of receptor R

$C$ - cell cluster

$i$ - cell group i

$j$ - cell group j

$M$ - the library-size normalized and log1p-transformed gene expression matrix

$X$ - normalized gene expression vector

We denote the two interaction proteins, via their genes $L$ & $R$, yet we use this for convenience as these can also denote the interaction of any other event category, such as those between membrane-bound or extracellular matrix proteins. Furthermore, in the case of heteromeric complexes $L$ & $R$ denote the summarized expression of the complex.

- **CellPhoneDBv2**

Magnitude**:** 1) $LRmean_{k,i,j} = \frac{L_{C_i} + R_{C_j}}{2}$

Specificity**:** A permutation approach also adapted by other methods, see 4)

- **Geometric Mean**

Magnitude: 2) $LRgeometric.mean_{k,i,j} = \sqrt{L_{C_i} \cdot R_{C_j}}$

Specificity**:** An adaptation of CellPhoneDB's permutation approach; see 4)

- **CellChat†**

Magnitude: 3) $$LRprob_{k,i,j} = \frac{TriMean(L_{C_i}) \cdot TriMean(R_{C_j})}{Kh + TriMean(L_{C_i}) \cdot TriMean(R_{C_j})}$$

where *Kh* = 0.5 by default and `TriMean` represents Tuckey's Trimean function:

$$TriMean(X) = \frac{Q_{0.25}(X) + 2 \cdot Q_{0.5}(X) + Q_{0.75}(X)}{4}$$

Specificity: An adaptation of CellPhoneDB's permutation approach; see 4)

† Note that the original CellChat implementation also uses information of mediator proteins and pathways, which is specific to the CellChat resource. Since we can use any resource with LIANA, we do not typically utilize this information, and hence the implementation of CellChat's scoring functions in LIANA was simplified to be resource-agnostic.

4) $$p\text{-value}_{k,i,j} = \frac{1}{P} \sum_{p=1}^{P} [fun_{permuted}(L^*_{C_i}, R^*_{C_j}) \geq fun_{observed}(L^*_{C_i}, R^*_{C_j})]$$

where $P$ is the number of permutations, and $L^*$ and $R^*$ are ligand and receptor expressions summarized according to the aggregation function per cluster used by each method, i.e. by default the arithmetic mean for CellPhoneDB and Geometric Mean, and TriMean for CellChat.

- **SingleCellSignalR**

Magnitude**:** 5) $$LRscore_{k,i,j} = \frac{\sqrt{L_{C_i} R_{C_j}}}{\sqrt{L_{C_i} R_{C_j}} + \mu}$$

where μ is the mean of the expression matrix $M$

- **NATMI**

Magnitude: 6) $$LRproduct_{k,i,j} = L_{C_i} R_{C_j}$$

Specificity: 7) $$SpecificityWeight_{k,i,j} = \frac{L_{C_i}}{\sum^n L_{C_i}} \cdot \frac{R_{C_j}}{\sum^n R_{C_j}}$$

- **Connectome**

Magnitude: 6) $$LRproduct_{k,i,j} = L_{C_i} R_{C_j}$$

Specificity: 8) $$LRz.mean_{k,i,j} = \frac{z_{L_{C_i}} + z_{R_{C_j}}}{2}$$

where $z$ is the z-score of the expression matrix $M$

- **Log2FC**

Specificity: 9) $$LRlog2FC_{k,i,j} = \frac{\text{Log2FC}_{C_i,L} + \text{Log2FC}_{C_j,R}}{2}$$

where log2FC for each gene is calculated as:

10) $$log2FC = \log_2(\text{mean}(X_i)) - \log_2(\text{mean}(X_{\text{not}_i}))$$

When generating a consensus from the different methods in LIANA, a rank aggregate is calculated for the *magnitude* and *specificity* scores from the methods separately. First, a normalized rank matrix[0,1] is generated separately for magnitude and specificity as:

11) $$r_{i,j} = \frac{rank_{i,j}}{\max(rank_i)} \quad (1 \le i \le m, 1 \le j \le n)$$

where $m$ is the number of ranked score vectors, $n$ is the length of each score vector (number of interactions), $rank_{i,j}$ is the rank of the j-th element (interaction) in the i-th score rank vector, and $max(rank_i)$is the maximum rank in the i-th rank vector.

For each normalized rank vector $r$, we then ask how probable it is to obtain $r^{null}_{(k)} <= r_{(k)}$, where $r^{null}_{(k)}$ is a rank vector generated under the null hypothesis. The RobustRankAggregate (https://github.com/cran/RobustRankAggreg) method expresses the probability $r^{null}_{(k)} <= r_{(k)}$ as $\beta_{k,n}(r)$ through a beta distribution. This entails that we obtain probabilities for each score vector $r$ as:

12) $$p(r) = \underset{1,\ldots,n}{min} \beta_{k,n}(r) * n$$

where we take the minimum probability ρ for each interaction across the score vectors, and we apply a Bonferroni correction to the P-values by multiplying them by $n$ to account for multiple testing.

13) LIANA considers interactions as occurring only if the ligand and receptor, and all of their subunits, are expressed in a certain proportion of the cells (0.1 by default) in both clusters involved in the interaction. This can be formulated as an indicator function as follows:

$$I\left\{L_{C_j}^{expr.prop} \ge 0.1 \text{ and } R_{C_j}^{expr.prop} \ge 0.1\right\}$$
